# Quantifying geographic accessibility to improve efficiency of entomological monitoring

**DOI:** 10.1101/779561

**Authors:** Joshua Longbottom, Ana Krause, Stephen J. Torr, Michelle C. Stanton

**Affiliations:** Department of Vector Biology, Liverpool School of Tropical Medicine, Liverpool, L3 5QA; Centre for Health Informatics, Computing and Statistics, Lancaster Medical School, Lancaster University, Lancaster, LA1 4YW

**Keywords:** Accessibility, Entomology, Remote sensing, Resistance, Vector control

## Abstract

**Background:** Vector-borne diseases are important causes of mortality and morbidity in humans and livestock, particularly for poorer communities and countries in the tropics. Large-scale programs against these diseases, for example malaria, dengue and African trypanosomiasis, include vector control, and assessing the impact of this intervention requires frequent and extensive monitoring of disease vector abundance. Such monitoring can be expensive, especially in the later stages of a successful program where numbers of vectors and cases are low.

**Methodology/Principal Findings:** We developed a system that allows the identification of monitoring sites where pre-intervention densities of vectors are predicted to be high, and travel cost to sites is low, highlighting the most efficient locations for longitudinal monitoring. Using remotely sensed imagery and an image classification algorithm, we mapped landscape resistance associated with on- and off-road travel for every gridded location (3m and 0.5m grid cells) within Koboko district, Uganda. We combine the accessibility surface with pre-existing estimates of tsetse abundance and propose a stratified sampling approach to determine the most efficient locations for longitudinal data collection. Our modelled predictions were validated against empirical measurements of travel-time and existing maps of road networks. We applied this approach in northern Uganda where a large-scale vector control program is being implemented to control human African trypanosomiasis, a neglected tropical disease (NTD) caused by trypanosomes transmitted by tsetse flies. Our accessibility surfaces indicate a high performance when compared to empirical data, with remote sensing identifying a further ~70% of roads than existing networks.

**Conclusions/Significance:** By integrating such estimates with predictions of tsetse abundance, we propose a methodology to determine the optimal placement of sentinel monitoring sites for evaluating control programme efficacy, moving from a nuanced, ad-hoc approach incorporating intuition, knowledge of vector ecology and local knowledge of geographic accessibility, to a reproducible, quantifiable one.

**Author Summary:** Assessing the impact of vector control programmes requires longitudinal measurements of the abundance of insect vectors within intervention areas. Such monitoring can be expensive, especially in the later stages of a successful program where numbers of vectors and cases of disease are low. Efficient monitoring involves a prior selection of monitoring sites that are easy to reach and produce rich information on vector abundance. Here, we used image classification and cost-distance algorithms to produce estimates of accessibility within Koboko district, Uganda, where vector control is contributing to the elimination of sleeping sickness, a neglected tropical disease (NTD). We combine an accessibility surface with pre-existing estimates of tsetse abundance and propose a stratified sampling approach to determine locations which are associated with low cost (lowest travel time) and potential for longitudinal data collection (high pre-intervention abundance). Our method could be adapted for use in the planning and monitoring of tsetse- and other vector-control programmes. By providing methods to ensure that vector control programmes operate at maximum efficiency, we can ensure that the limited funding associated with some of these NTDs has the largest impact.

## Introduction

Vector-borne diseases (VBDs) are important causes of mortality and morbidity in humans and livestock, particularly for poorer communities and countries in the tropics, accounting for an estimated 17% of the global burden of all infectious diseases (1). The control of VBDs, or their elimination as a public health problem, is dependent upon effective vector management, which includes pre-intervention surveys and subsequent longitudinal monitoring of vector abundance to assess the effectiveness of an intervention. Such monitoring is an important component of the overall costs of control.

To improve the efficiency of vector control programs, there is a requirement to identify optimal locations for longitudinal monitoring site placement. Ideally, these sites should be in locations that maximise information on the distribution and density of vectors while minimising costs of obtaining these data. In practice, most vector surveillance is opportunistic and lacks a rigorous framework (2). A more rational method would involve combining information on vector abundance with estimates of geographical accessibility, to identify sites across operational areas where pre-intervention catches are high and sampling costs are low. Towards this goal, we examined the utility of remotely sensed (RS) data to produce contemporary estimates of geographic accessibility to entomological sampling sites, using sleeping sickness control as an example application.

### Sleeping sickness control as an example application

Human African trypanosomiasis (HAT) is a neglected tropical disease (NTD) affecting remote areas of sub-Saharan Africa. The disease, also termed ‘sleeping sickness’, is caused by the protozoan parasite *Trypanosoma brucei* with two sub-species, *T.b.gambiense* and *T.b.rhodesiense*, causing Gambian (gHAT) and Rhodesian (rHAT) human African trypanosomiasis respectively. The burden of the Gambian form of the disease, for which humans are the main hosts, is >10 times that of the Rhodesian form, with annual reported cases being in the region of 2-3,000 (3). The World Health Organization (WHO) has targeted the elimination of gHAT as a “public health problem” by 2020, which is defined as a 90% reduction in areas reporting >1 case in 10 000 compared to 2000–2004, and <2000 annually reported cases globally (4). Several countries appear to be on track to achieve this target (5). Uganda is unique in that it is the only country where both gHAT and rHAT occur, albeit within different local level zones (6, 7). Vector control forms an important part of Uganda’s efforts against both forms of HAT (8, 9).

The important vectors of gHAT are Palpalis-group species of tsetse, which concentrate in riverine vegetation where, consequently, interventions are focused. In Uganda, tsetse control is being achieved through the deployment of Tiny Targets, small (20 × 50 cm) panels of insecticide-treated material which are deployed at 50-100m intervals along rivers (9, 10). Prior work produced estimates of tsetse abundance across Northern Uganda, identifying locations of high pre-intervention abundance (11), which has informed the identification of operational control areas.

Methods to quantify accessibility largely involve cost-distance analyses, which have been widely used within the field of public health in analyses mapping accessibility to healthcare (12–15). Such analyses require an input surface of landscape friction (‘resistance’) – estimates of associated travel cost for gridded cells within a Cartesian plane. The cost-distance analysis identifies the cumulative cost of traversing each cell based on the given resistance surface and an origin location – opting to traverse through cells associated with the lowest resistance values. The use of accessibility mapping in the planning and implementation of control programmes for vector-borne disease is novel and has the potential to improve the efficiency of monitoring VBD interventions.

In this paper, we use remotely sensed (RS) satellite data to derive a contemporary road network within Koboko district, Northern Uganda, where an existing tsetse control programme is in operation. To obtain a road network within this district, we compare the utility of RS data at two differing spatial resolutions (one source characterising locations within the district as 3 × 3m grid cells on a Cartesian plane, and another as 0.5 × 0.5m grid cells) (16, 17), and an existing open source dataset detailing road locations (18). Image classification algorithms, specifically maximum likelihood estimators were used to detect dirt and tarmac roads within the RS imagery (19). Ground truth tracking (GPS) data detailing motorbike speeds along roads within the district were used to assign on-road travel costs to each grid cell. We used published estimates of time taken to traverse through different densities of vegetation to assign resistance values to off-road grid cells (20, 21). Resistance surfaces were validated using withheld ground-truth tracking data, comparing observed and predicted travel times within a linear regression. The resulting resistance surfaces were used within a least-cost path algorithm to identify cumulative costs to locations of high tsetse abundance (11). We apply a stratified sampling approach to determine locations which are associated with low cost (lowest travel time) and potential for rich longitudinal data collection (high pre-intervention abundance).

Here, by combining field data on travel time along varying road types and remotely sensed imagery, we describe the process of producing a high-resolution accessibility surface. By integrating such estimates with predictions of tsetse abundance, we propose a methodology to determine the optimal placement of sentinel monitoring sites for evaluating the efficacy of a tsetse control programme, moving from a nuanced, ad-hoc approach incorporating intuition, knowledge of vector ecology and local knowledge of geographic accessibility to a reproducible, quantifiable one. The work described here is presented in the context of tsetse control, but the methods used are applicable to a wide range of vector-borne diseases.

## Materials and Methods

### Study area

The focal area of this study was Koboko District, located within the West Nile Region of Uganda. The West Nile region consists of eight districts, with current and planned intervention initiatives (i.e. the Tiny Target programme), operating in seven. Koboko district covers roughly 860km^2^ and has a population of 229,200 people (22). Between 2000 and 2018, 14.6% (620/4235) of gHAT cases reported from Uganda occurred in Koboko, but the incidence of gHAT is in decline as a consequence of an integrated programme of screening and treatment of the human population and, more recently, vector control (23). A map showing the location of existing, and planned intervention areas within West Nile Region is provided as Fig. S1, highlighting the position of Koboko within these intervention districts.

### Field methodology and data collection

To obtain data informing variation in speeds along road class, technicians making routine visits to traps within Koboko were provided with GPS devices. The recording of GPS tracks was performed during three time periods in the dry season: May-June 2017, February-April 2018, and December 2018-January 2019. Trap attendants within Koboko operate using motorbikes; therefore, observed speeds were representative of motorbike-based travel. Devices were configured to record track points at ~15-second intervals.

### Obtaining remotely sensed satellite data

To compare the effect of different spatial resolutions of satellite data on the ability to identify roads, we used two differing sources of remotely sensed imagery. Imagery obtained from PlanetScope™ satellites, captured on February 12^th^, 2018 were utilised. PlanetScope™ imagery is provided at a 3m × 3m resolution, and includes the following four spectral bands: blue (455 – 515 nm), green (500 – 590 nm), red (455 – 515 nm), and near infrared (780 – 860 nm) (16, 24). PlanetScope™ data are freely accessibly through an education and research program account. Data captured through the Pléiades-1A satellite, available at a 0.5m × 0.5m resolution and captured on 27^th^ December 2016 were used to represent high-spatial resolution imagery (25). Imagery captured on this date was the most contemporary data available. The Pléiades-1A imagery similarly consists of the same four spectral bands as PlanetScope™. Data obtained by Pléiades-1A is available by request through Airbus (previously known as the European Aeronautic Defence and Space Company) (17).

### GPS data review and cleaning

To calculate travel speeds, the time-difference between subsequent points within a track and the Euclidean distance between these points were used within the following formula (Equation 1):

Where *x*_*i*_ represents the GPS coordinate of point *i*, *t*_*i*_ represents the time recorded for point *i* and ∥·∥ represents the Euclidean distance:

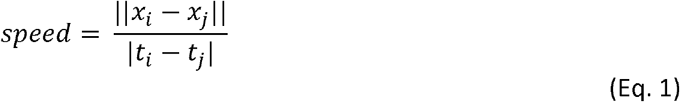

Recorded points with a speed <1km/hr were assumed to be stationary points (based on average walking speeds (26)), and were removed from the track dataset. Similarly, we removed data points for which the speed exceeded 150 km/hr (93.2 mph) as these were likely to be artefacts created due to errors with location positioning and are not representative of true travel speed.

### Open street map validation

To determine the accuracy of currently available open source data, OpenStreetMap (OSM) geolocated roads, and roads visible within 0.5m and 3m satellite data were compared. Shapefiles detailing mapped roads hosted by OSM were retrieved from Geofabrik OSM Data Extracts on March 3^rd^, 2018, to align with the dates during which field-obtained tracking data were collected (18). A 1 km × 1km fishnet constructed for Koboko district was used to produce a random sample of 25 grid squares for manual digitisation. The digitisation process consisted of tracing over visible roads and tracks, as seen in the 0.5m resolution imagery (metric one), or as seen in the 3m resolution imagery (metric two). The length of digitized road obtained from each of the three sources was calculated in metres.

### Remote sensing image preparation

In total, 14 scenes covering an area of 745.8 km^2^ were downloaded from Planet.com. To produce one complete surface, overlapping scenes were merged using ArcGIS (version 10.4), and the composite image was cropped to district boundaries. Imagery obtained from Pléiades-1A (0.5m) were provided as a pre-prepared mosaic.

### Image classification

To aid image classification, image segmentation utilising a mean-shift approach was first performed within ArcGIS (version 10.4). Mean-shift segmentation is a process that identifies segments in imagery by grouping adjacent pixels that have similar spectral characteristics; a detailed introduction and theory related to mean-shift segmentation algorithms can be found within Demirović 2019 (27). We utilised the “*Segment Mean Shift*” tool within the “*Spatial Analyst Toolbox*” in ArcGIS, with the following default parameters: spectral detail = 15.5, spatial detail = 15, minimum segment size (in pixels) = 20. Following mean-shift segmentation, we applied a maximum likelihood (ML) classification algorithm using an equal *a priori* probability weighting to identify the class in which each cell had the highest probability of being a member. The ML classification algorithm considers both the variances and covariances of pixels assigned to ‘classes’ (groups of pixels relating to a specific type of land-cover, in this instance), selected within a signature training file (19). Under the assumption that the distribution of a class sample is normal, each class was characterized by the mean vector and the covariance matrix. Given these characteristics, for each cell value within the remotely sensed imagery, the statistical probability of a cell belonging to each class is calculated and an appropriate classification is assigned (19). We opted to use the following classes within this analysis: dirt road and/or track, tarmac road, dense vegetation (for example: woodlands, forest, bushwood and shrubwood), grassland (for example: grassland, meadow, steppe and savannah) and barren land. Signature files for use in the ML classification were produced by manually tracing and assigning pixels within the remotely sensed imagery to one of the five classes described above. Classification was performed using the “*Train Maximum Likelihood Classifier*” tool within the “*Spatial Analyst Toolbox*” in ArcGIS. To account for “salt and pepper” speckling effects representative of potentially misclassified and/or isolated cells, we performed post-classification processing. This processing stage included filtering to remove isolated cells (28), smoothing to smooth rugged class boundaries (29), and generalizing to reclassify small regions of isolated cells (30). Post-classification cleaning was performed in ArcGIS.

### Classification validation

A total of 500 accuracy assessment points were randomly generated for each classified surface (i.e. 3m × 3m and 0.5m × 0.5m imagery). A step-by-step comparison was then made for each randomly selected point, noting the algorithm-derived class and the manually assigned (ground-truth) class. Utilising this information, a confusion matrix was constructed for each image source. Accuracy was calculated with respect to both omission and commission rates, where omission refers to instances where a feature (point) is omitted from the evaluated category, and commission refers to instances where a feature is incorrectly assigned to the category being evaluated.

### Road network update

Using the outputs from the image classification process, the GPS tracking data, and available OSM data, two contemporary road networks (one per remotely sensed data source) were produced. Cleaned, field-obtained tracking points were used to inform estimates of average travel speeds along selected roads as follows. Tracking points were converted to polylines, consisting of line segments constructed from five trailing points. These segments were assigned a mean observed speed by calculating the Euclidean distance of each segment and incorporating start and end times. These segments were then rasterised, resulting cells were stacked, and overlapping cells resulting from replicate trips across all tracking days were averaged. This produced a surface indicating the average observed speed for each cell. Tracks obtained during December 2018 were withheld from this network and were used for validation (see below). A surface detailing urban and rural locations (31) was used to categorise roads as being within urban or rural areas. This classification was paired with the Ugandan Traffic and Road Safety Act detailing maximum speed limits based on roads within urban/built-up areas and rural areas. Characterising roads by these features imply a legal maximum speed for each road representative of true travel speeds. Classified urban and classified rural cells were assigned the speeds given in Table 1, as informed by the official Traffic and Road Safety Act 2004 (32) and the Highway code (33).

**Table 1.** Assigned travel speeds to roads lacking ground-obtained tracking data.

| Road type | Speed (km/h) |  |
| --- | --- | --- |
|  | Built-up area | Rural area |
| Paved | 50 | 100 |
| Gravel/dirt | 50 | 80 |

### Normalized Difference Vegetation Index analysis

As the majority of mapped roads do not lead directly to a river or tributary, trap attendants are required to traverse off-road in order to reach suitable habitats for trap placement. We therefore aimed to characterise the cost associated with off-road travel within our analysis. Utilising the two differing imagery sources, two separate NDVI surfaces were generated (Equation 2). During the NDVI calculation, output values were normalised to range between −1.0 and 1.0, representing greenness. Generally, output NDVI values ≤0 represent waterbodies including lakes and major rivers; values between 0.1 and 0.2 represent barren land, including areas of rock, sand, or snow; values between 0.2 and 0.3 represent shrub and grassland (areas of moderate vegetation), and values between 0.3 and 0.8 represent areas of dense vegetation (for example temperate and tropical rainforest) (34, 35).

Where *NIR* represents the near infrared band, and *R* represents the red band within the RS imagery:

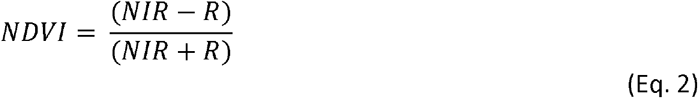

### Assigning off-road resistance values

Resistance values are values associated with a specific cost to traverse through a cell (time, in seconds). For this study, off-road resistance values were assigned utilising the NDVI outputs, with cost values ranging based on indicative terrain. Locations which contain dense vegetation are generally slower to navigate and therefore cells representative of these areas were associated with a higher resistance value; conversely, cells which represent areas with little to no vegetation were presumed to be easier to traverse and were assigned a lower resistance value. Average off-road walking speeds for differing terrains were obtained from published literature (20, 21) (Table 2).

**Table 2.** Resistance values (cell crossing time) associated with off-road travel.

| NDVI value | Off road walking speed (km/h) | Off road walking speed (m/s) | Cell crossing time ($t_i$) | |
| --- | --- | --- | --- | --- |
|  |  |  | 0.5 × 0.5m | 3m × 3m |
| $\leq 0$ | Essentially impassable | Essentially impassable | 200 | 200 |
| 0.1-0.2 | 3.5 | 0.97 | 0.73 | 4.37 |
| 0.2-0.3 | 2.48 | 0.69 | 1.03 | 6.14 |
| 0.3-0.8 | 1.49 | 0.41 | 1.73 | 10.34 |

### Resistance surface and cost-distance analysis

The updated road networks, featuring a cell crossing time based on assigned speeds (representative of on-road resistance), were combined with their respective NDVI resistance surface. To validate the generated surfaces, we used field-obtained tracking data (obtained December 2018) withheld from the road network construction. Sixty-three segments along the withheld tracks were used to create validation points. Using the resistance surface, the travel time from the start to the end point of each segment was generated utilising a least-cost path algorithm within QGIS 3.4.4 (36), plugin “*Least-Cost Path*” (produced by FlowMap Group (37)). The specific algorithm implemented is referred to as Dijkstra’s algorithm, and is an approach utilising graph theory to identify the shortest path between two nodes; the algorithm is described in detail in Dijkstra 1959 (38). A linear regression model was then fitted to the observed travel time data with predicted travel time being included as the only covariate to quantify the relationship between the two measures. The ability of the predicted travel time to each validation point to accurately predict the observed travel time was used to detect an association between the two, and to provide a means of adjusting the generated surface values if necessary. The accuracy of each resistance surface was defined by the coefficient p-values, and by root-mean-square error (RMSE). Utilising these resistance surfaces, two separate cost-distance analyses were performed (one per spatial resolution), each using the location of our district entomologist’s base as the origin. The cost-distance analysis again implemented Dijkstra’s algorithm, calculating the cumulative cost of travel from the origin to each grid cell in the resistance surface.

### Identifying optimal sentinel site placement

We performed a spatially stratified sampling approach to aid the identification of 104 least-cost, high abundance locations per 25km^2^ for sentinel site placement. Firstly, we produced a fishnet consisting of 5 km × 5 km grid squares across Koboko district, and assigned each grid square a sequential stratum identification number (see Fig. S2 for strata distribution). For each strata within the proposed intervention area, we ranked each cell by their predicted tsetse abundance values (11), and by their predicted travel time from the origin – as obtained from the cost-distance output. To account for spatial clustering, and to ensure a more even spatial distribution of sentinel sites, we retained the cell with the highest predicted abundance and lowest associated cost per 50m × 50m area. We calculated the cumulative rank for each cell within the de-clustered dataset, where predicted abundance values were ranked from high to low, and accessibility values ranked from low to high. We retained two locations (paired sites) with the lowest cumulative rank per sampled strata, with these locations being identified as the optimal placement for sentinel monitoring sites.

### Utilising the travelling salesperson problem (TSP) to identify the optimal route

Once the optimal location of monitoring sites was identified, we applied the travelling salesperson problem (TSP) to identify the most efficient order in which to visit each site. The TSP is an optimisation problem in which the following question is addressed: “Given a list of cities and distances between each pair of cities, what is the shortest possible route that visits each city and returns to the origin city?”(39). We adapt this problem to answer “Given a list of monitoring locations and travel times between each pair of locations, what is the shortest possible route that visits each monitoring site and returns to the origin location?”. We solve this through the implementation of “Concorde’s algorithm” (39), through the TSP package in R (40). First, the 3 × 3m friction surface was converted into a transition matrix through use of the “*transition*” function in the gdistance R package (41). Second, the pairwise distances between each site was calculated to produce a distance matrix, through use of the “*costDistance*” function in the gdistance package. We then implemented the TSP using the function “*TSP*” and the distance matrix, and solved the TSP with “*solve_TSP*”; both functions are from the TSP R package. By following the route identified by solving the TSP, and incorporating 30-minute stays at each pair of sites to deploy traps and/or collect samples, we group sites into ‘clusters’ which are feasible to visit within a 5-hour sampling day.

## Results

### GPS data collection

To inform estimates of on-road travel cost for each 3m × 3m and 0.5m × 0.5m cell within Koboko district, Northern Uganda, we obtained tracking data during three periods: May-June 2017, February-April 2018, and December 2018-January 2019. Tracks collected between May 2017 - April 2018 were used to inform road speeds, and tracks collected between December 2018-January 2019 were withheld for validating the resistance surfaces (Fig. S3).

### OpenStreetMap accuracy assessment

Analyses evaluating the accuracy of an existing, community-driven, open-source road network (from OpenStreetMap), indicate that at least one road exists within the OpenStreetMap (OSM) dataset for 17 out of 25 randomly sampled 1km^2^ grid squares across Koboko district (mean road length = 1.97 km). Only one out of 25 grid squares contained no visible roads across sources (i.e. 0.5m imagery, 3m imagery, and OSM). When comparing total road length visible in 3 × 3m imagery with that charted by OSM, the two sources show close agreement (97.43% similarity [total road length across 25km^2^], paired t-Test *p* = 0.91), however, when comparing the 0.5 × 0.5m imagery and the OSM dataset, only 28.16% of digitised roads are charted by OSM (paired t-Test *p* < 0.001, Fig. 1, Table S1, Fig. S4). This section of the analysis provided the rationale for the classification of 0.5m imagery, with the inclusion potentially capturing up to 71% more roads than OSM within the study area.

**Fig. 1.**
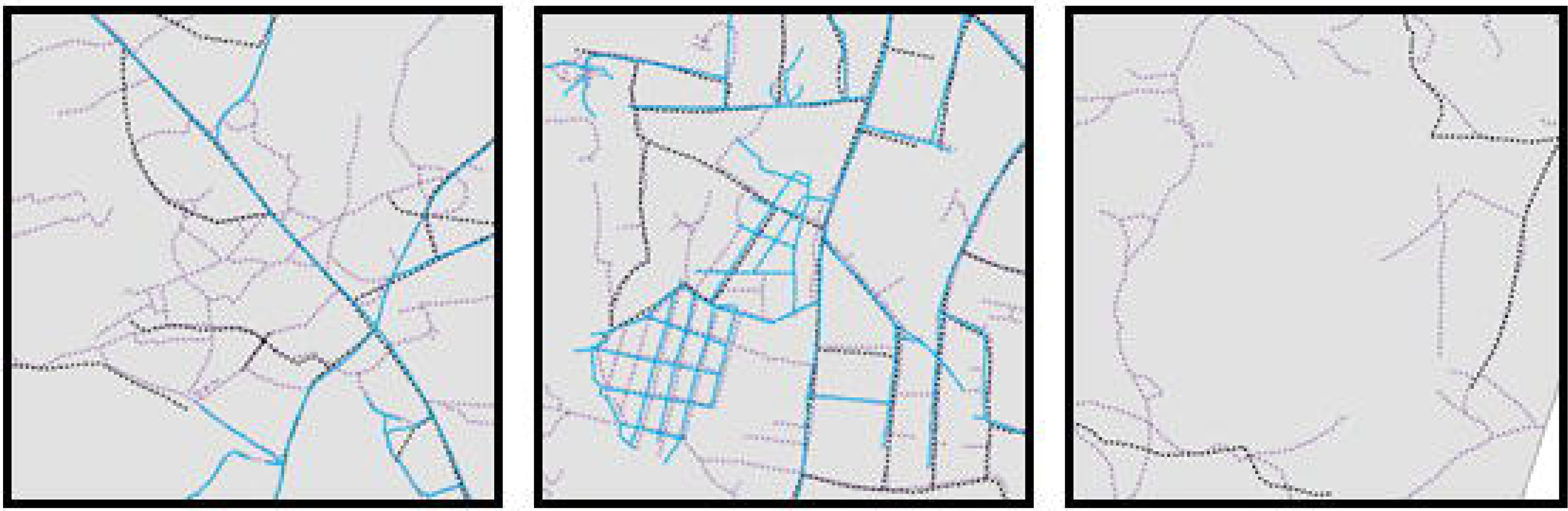
Example of composite images of digitised road networks within Koboko district. Purple roads represent roads visible in 0.5m imagery (17), as digitised in this study; black roads represent roads visible in 3m imagery (24), as digitised in this study, and light blue roads represent roads available within the OSM dataset (18). The overlap of all three colours indicate areas of consistency across sources.

### Image classification

Classification of two differing sources of remotely sensed imagery (0.5 × 0.5m and 3 × 3m) yielded varying accuracies across classes, and across spatial resolutions, with accuracy values ranging from 38% to 89% for dirt roads and 5% to 84% for tarmac roads for 3m and 0.5m imagery respectively (Table 3; Fig. 2). Overall image classification accuracy, considering all five classes utilised (dirt road and/or track, tarmac road, dense vegetation, grassland and barren land), ranged from 53% (3m) to 78% (0.5m), with 0.5m imagery proving to be more effective at identifying both dirt and tarmac roads than the 3m imagery.

**Table 3.** Maximum likelihood classification (MLC) accuracy assessment validation values for each class. Values represent the percentage of correctly classified cells (classified *vs* ground truth) for the five classes of interest.

| Class | 3m imagery |  |  | 0.5m imagery |  |  |
| --- | --- | --- | --- | --- | --- | --- |
|  | Correct | Incorrect | Accuracy (%) | Correct | Incorrect | Accuracy (%) |
| Dirt road and/or track | 38 | 62 | 38.00 | 97 | 11 | 89.81 |
| Tarmac road | 5 | 95 | 5.00 | 84 | 16 | 84.00 |
| Dense vegetation | 91 | 9 | 91.00 | 95 | 5 | 95.00 |
| Grassland | 65 | 34 | 65.65 | 80 | 20 | 80.00 |
| Barren land | 70 | 30 | 70.00 | 38 | 54 | 41.30 |
| **Overall** | **269** | **230** | **53.91** | **394** | **106** | **78.80** |

**Fig. 2.**
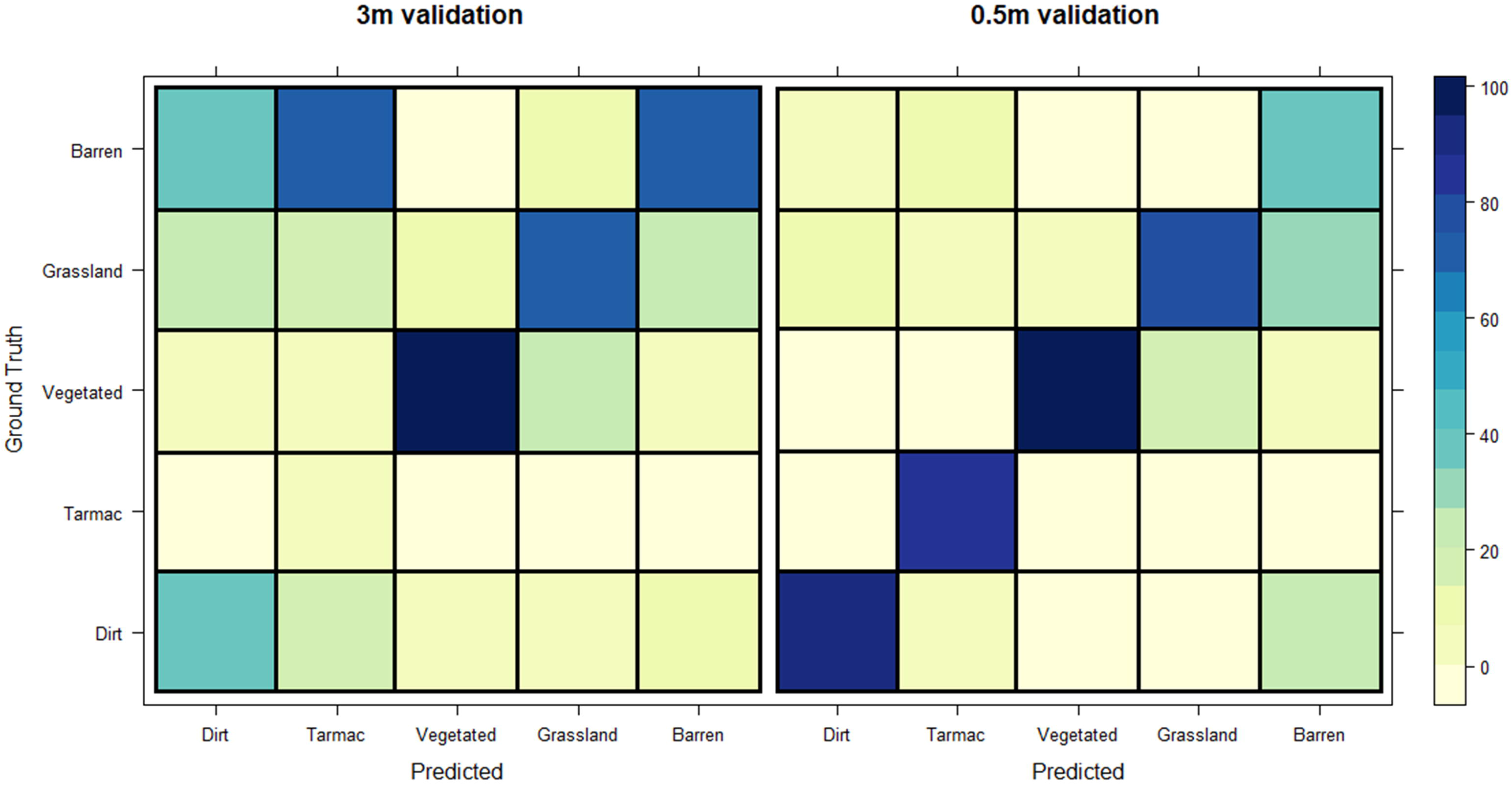
Confusion matrices for the classification of each surface (Left: 3m, Right: 0.5m). Diagonal squares (bottom left to top right) indicate the percentage of correctly classified cells per class.

### Resistance surface and cost-distance analysis

The accuracy of the resistance surfaces was assessed by investigating the relationship between observed travel times and predicted travel times using withheld field-obtained GPS tracks and a linear regression. Predicted values produced utilising the 3m resistance surface have a much closer alignment with ground truth (observed) values, root-mean-square error (RMSE) = 3.93 (3m) than the 0.5m resistance surface (RMSE = 6.01). In separate regressions with validation data from both surfaces, we identify that there is a significant association between observed and predicted values (*p* < 0.001 (0.5m) and *p* < 0.001 (3m)), indicating a high performance of each surface, with the 3m surface showing a stronger relationship with less variability (*R*^2^ = 0.66 vs *R*^2^ = 0.49, 3m and 0.5m respectively). Summaries of resistance surface validation are provided within Fig. S5 and Table 4. Output cost-distance surfaces detailing the travel time from the location of our field station to each gridded cell within Koboko district are provided as Fig.3.

**Table 4.** Model summaries for resistance surface validation. Summary statistics from four separate linear regressions are provided.

|  | 3m resistance surface |  | 0.5m resistance surface |  |
| --- | --- | --- | --- | --- |
|  | Training data | Validation data | Training data | Validation data |
| p-value | < 0.001 | < 0.001 | < 0.001 | < 0.001 |
| Coefficient | 1.10 | 0.59 | 0.78 | 0.34 |
| RMSE | 1.11 | 3.93 | 0.85 | 6.01 |
| $R^2$ | 0.93 | 0.66 | 0.95 | 0.49 |

**Fig. 3.**
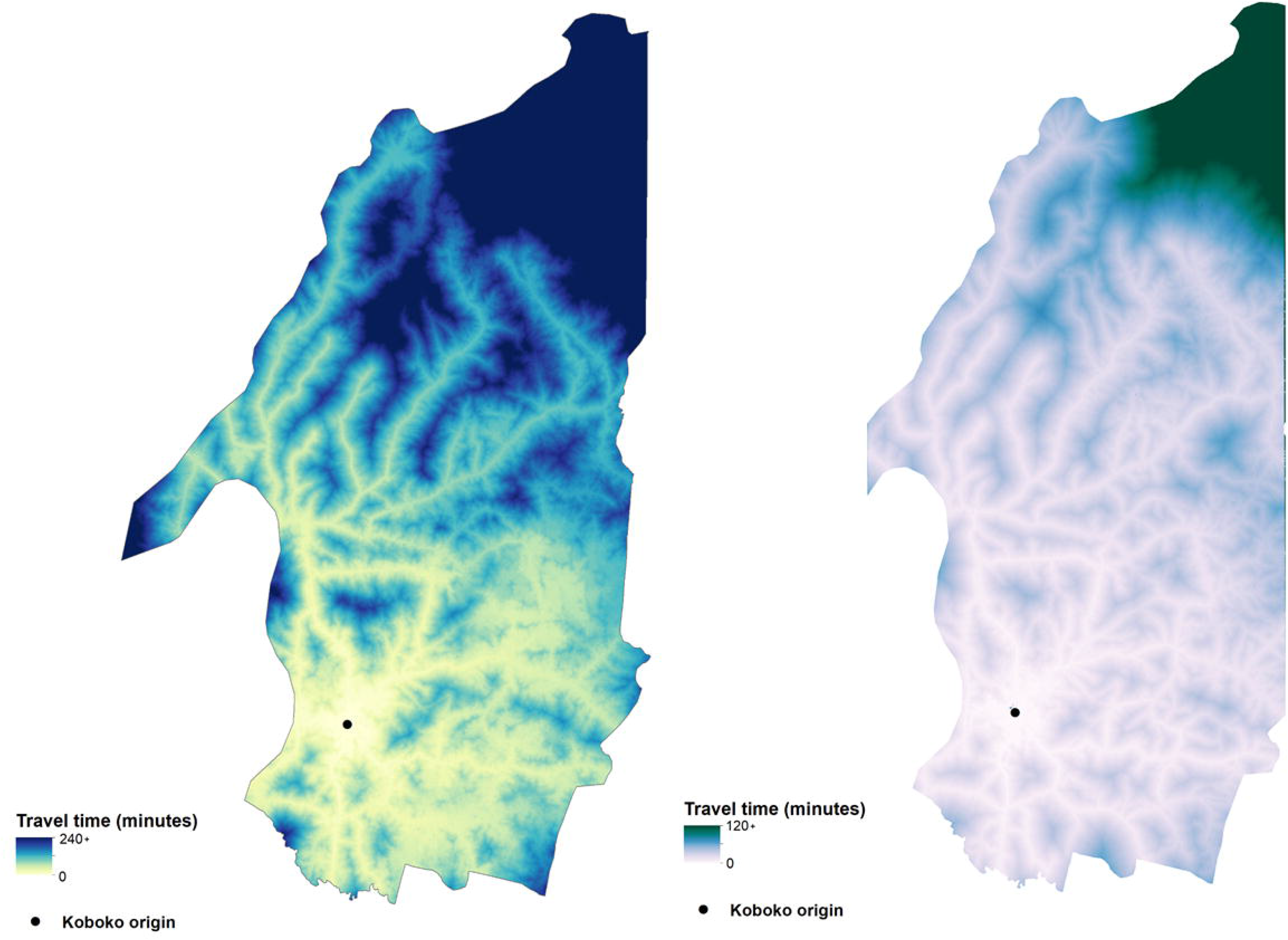
Cost-distance surfaces. Figures show the cumulative travel time from the field site origin (black point), to each subsequent cell within the surface. Left: 3m cost-distance surface, Right: 0.5m cost-distance surface. This figure was generated using ArcGIS version 10.4 (42), and products derived in this study from image classification of Planet (3m)(24) and Airbus (0.5m)(17) satellite imagery.

### Identification of optimal sentinel site placement

Utilising the 3m cost-distance surface and a predictive surface of tsetse abundance (11), we identified the optimal placement of 104 sentinel sites within the current intervention area (52 paired locations) (Fig. 4). Such sites are positioned within the most easily accessible, high abundant locations for 26 unique 5 × 5 km strata across the intervention area. Optimal sentinel-site placement identifies locations with abundance values ranging from 0.04– 19.57 (mean = 5.21) flies per cell, and locations which are within 5.55 - 151.81 (mean = 68.42) minutes from the field station location.

**Fig. 4.**
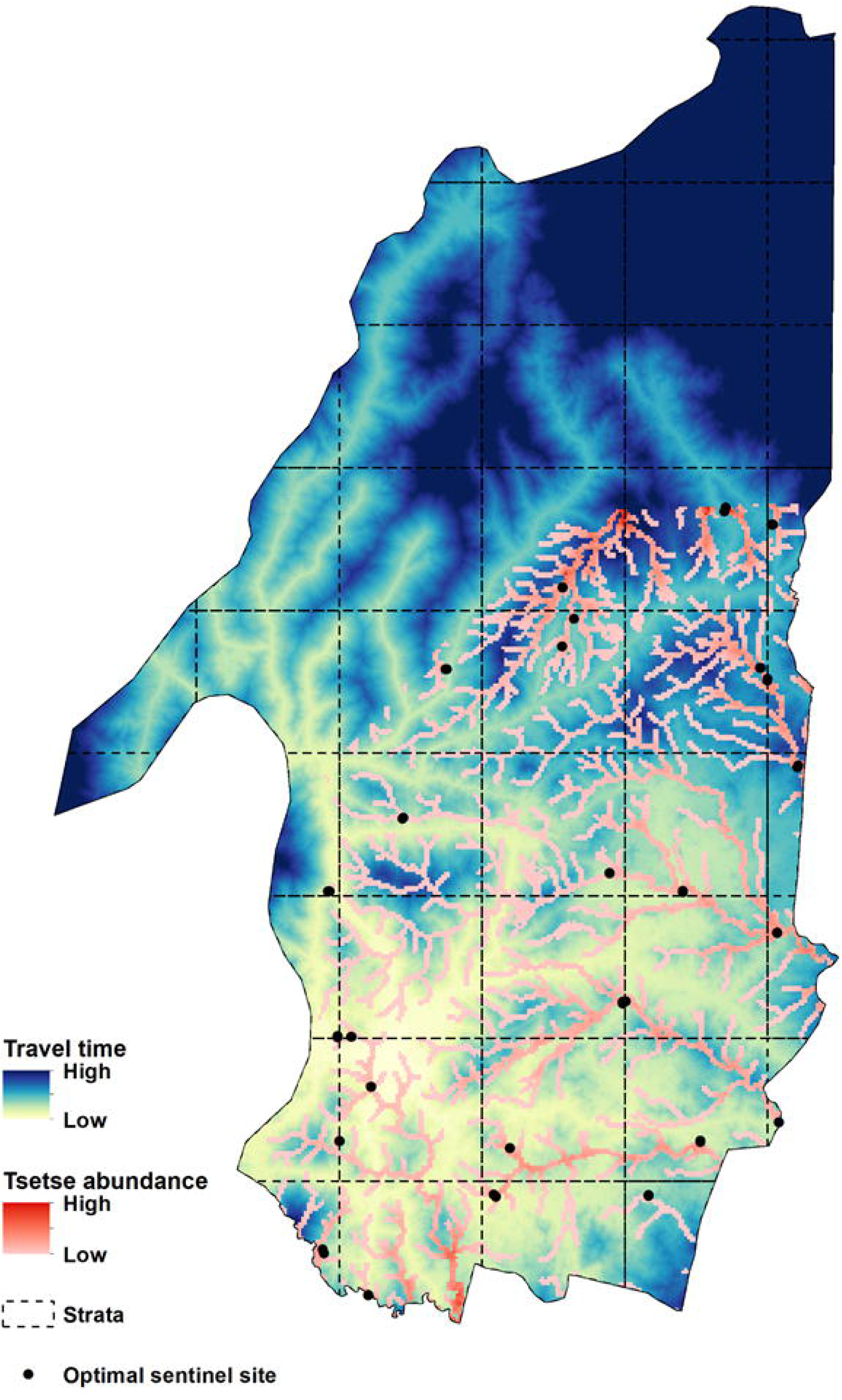
Optimal placement of sentinel sites (max two sites per grid square [25km^2^]) within Koboko district. Location of optimal sites visualised alongside the 3m accessibility surface (this study) and tsetse abundance surface (11), dashed lines represent the 5 × 5km sampling strata used to allocate optimal sites. This figure was generated using ArcGIS version 10.4 (42).

### Identification of the optimal route

Utilising the coordinates of the 52 paired monitoring site locations, derived above, we implemented the traveling salesperson problem (TSP) to identify the optimal route in which to visit these sites. The result of the TSP is shown as Fig. 5. Based on the assumption that the field-team will spend up to 5 hours sampling per day, incorporating travel times, we grouped sites to identify sampling clusters to visit per day. We show that a sampling period of four days is required to ensure that all sample locations are visited.

**Fig. 5.**
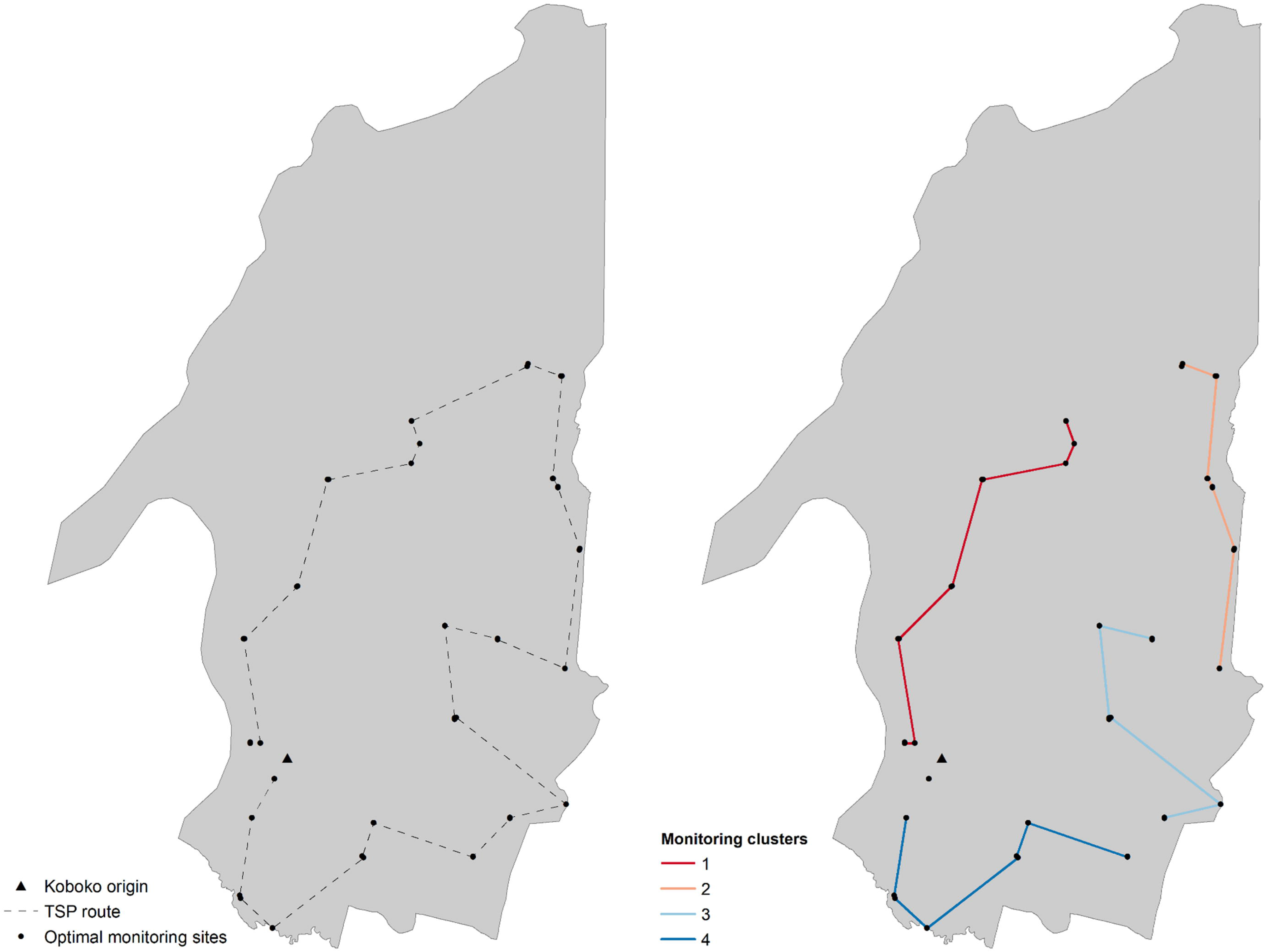
Left: Optimal route in which to sample the identified sentinel sites within Koboko district. Right: Clusters of sentinel sites are identified by enforcing a maximum sampling and travel period of 5-hours within Koboko district. This figure was generated using ArcGIS version 10.4 (42).

## Discussion

This analysis investigated the ability of high-resolution satellite imagery to inform estimates of accessibility to entomological sampling sites, using tsetse control as an example application. We started by scrutinising the completeness of an existing open source road network for Koboko district, Uganda, comparing charted roads with those obtainable from manual digitisation of remotely sensed (RS) imagery at two differing spatial resolutions. Results from this section of the analysis indicate that, for this region of Uganda, roads visible within 3m imagery matched 97.43% of roads identified in OpenStreetMap (OSM) (paired t-Test *p* = 0.91) (Fig. 1, Table S1). Comparing roads visible within 0.5m RS imagery, and those charted by OSM, yields 28.16% consistency across sources (paired t-Test *p* < 0.001) (Table S1).

As data published on OSM is the result of community contributions incorporating local knowledge, data coverage is often inconsistent. The recent establishment of several refugee camps across the West Nile Region has resulted in increased road mapping efforts within this area, which explains the high levels of coverage seen here (43). OpenStreetMap completeness varies globally and the analyses we have developed will be particularly useful in places where OSM and standard sources of information on road networks are scant (44).

Part of our analysis aimed to infer the effect of including spatially disaggregated data on estimates of accessibility, detailing whether the extra information obtainable from 0.5m imagery produces refined estimates. The results of a maximum likelihood classification algorithm indicate a high ability to identify roads and associated features within the 0.5m imagery, mirroring that seen by manual digitisation (Table 3; Fig. 2). Results from image classification also indicate that the spatial detail available within 3m imagery is too coarse to classify roads in this district accurately (38% and 5% accuracy for dirt and tarmac roads respectively). This result is to be expected as the majority of roads within Koboko district rarely exceed a width of 3m, resulting in decreased visibility; narrow roads are likely to be common across large parts of rural Africa (45). The utility of 3m imagery may be greater in more developed areas, where roads exceed 3m in width.

Despite a higher image classification accuracy and a better model fit to training data, the 0.5m resistance surface appears to under-perform when presented with withheld GPS tracking data compared to the 3m resistance surface (Table 4, Fig. S5). Both resistance surfaces show a significant linear relationship between observed and predicted values, however, the 3m resistance surface has a lower root-mean-square error (3.93 *vs* 6.01 respectively). This under-performance may be due to the increased number of roads within the 0.5m resistance surface, and some of the assumptions made regarding travel along roads of differing class. While we have used the best possible information available to us, there will invariably be additional factors that may affect how accessible a location is. Should, in practice, a location be more difficult to access than predicted using our approach, an alternative location will be selected based both on the outcome of this approach and field-based information. We envisage this process to be somewhat iterative, with new GPS data collected during the first visit to a proposed monitoring site. This new data may be used to improve surface validation and refine some of the assumptions made during the approach described here. When using the surfaces to identify optimal placement of sentinel-sites, the relative travel-time to each cell is as informative as the actual travel-time. Despite varying RMSEs, the significant relationship between predicted and observed travel times, support the utility of the generated surfaces.

By combining the generated 3m accessibility surface (Fig. 3) with previously published estimates of tsetse-abundance (11), we provide a novel framework for the identification of efficient locations in which to place sentinel-monitoring sites (Fig. 4). Previous methods to inform the placement of sentinel-monitoring sites have been based on intuition, incorporating knowledge of tsetse ecology and local knowledge of roads within an intervention area. Here, we further quantify this process, providing a more robust approach that can be applied to a range of vector-borne diseases. The movement from a nuanced, ad-hoc process to an evidence-based one will allow for a more efficient assessment of tsetse control programmes. Although we have provided a quantifiable approach for prioritising spatial sampling of disease vectors, we are aware that knowledge of additional country and context specific factors such as varying vector behaviours and geographic accessibility are invaluable for designing and implementing an effective monitoring program. Such approaches should be tailored for the vector, disease, and country of interest, with the work described here providing a framework from which to build. Local knowledge can still be useful in the design and implementation of this approach, potentially when identifying changes in accessibility (such as the creation or disuse of roads), or through refinement of selected sites. The application of the methods used here to the context of intervention monitoring and assessment is novel, and the refinement of results has several cost-effective implications as vector control expands to other areas within the region.

The distribution and abundance of disease vectors dynamically change in response to variations in biotic and abiotic conditions (46, 47). The methodology described here is receptive to new surfaces detailing expanding or decreasing species ranges, however our approach is focused on identifying static monitoring sites based off conditions at the time of implementation of the intervention. Periodic updates to OSM data may be used to generate contemporary geographic accessibility surfaces reflecting the creation of new road networks or the disuse of others. Although this methodology has the potential for dynamic updates, we are aware that our approach requires a technical understanding of GIS and remote sensing, factors which may prevent uptake and application in developing countries outside of the framework of internationally supported programs. These factors may be addressed through capacity strengthening programmes, where GIS skills can be integrated as part of the curriculum.

Several important vector-borne NTDs have been targeted for elimination as a public-health problem by 2020 within the WHO NTD roadmap (4). Unfortunately, however, the burden of numerous VBDs will continue beyond the ambitious 2020 target (48–50). As evident within the WHO roadmap, both disease and vector surveillance form large components of most elimination strategies; however, the Strategic and Technical Advisory Group (STAG) for NTDs also recognise the need for a better understanding of the economic aspects of NTD control. By providing methods to ensure that vector control programmes operate at maximum efficiency, we can ensure that the limited funding associated with some of these NTDs has the largest impact.

Although this analysis does not serve as an economic evaluation of methods to assess control programme efficacy, previous work has shown that vehicle running and travel costs are within the top five associated costs of running a tsetse control programme (51, 52), with staff salaries being the most expensive element. By strategically placing sentinel-monitoring sites in locations that are associated with a low accessibility cost, programmes can reduce costs associated with travel (e.g., fuel, maintenance) and staff expenses, with current costs of tsetse monitoring being ~9.0$/km^2^/year (10.6% of tsetse control programme budgets) (52). The accessibility surface may also contribute toward cost-effective planning of pre-intervention surveys, which are responsible for roughly 6% of control program budgets (52). Furthermore, by informing the positioning of these sites by additional metrics, such as pre-intervention abundance, we identify locations that may provide more accurate evaluations of control efficacy. Further research should be performed to evaluate the precise economic gains of this approach.

Accessibility, in general, is a very sought-after metric and the methodology applied here, although currently restricted to one district in Northern Uganda and limited to the purpose of identifying accessible tsetse monitoring sites, could inform other accessibility analyses within the area such as access to HAT diagnostic centres, and may be applied to a range of vector-borne diseases.

## Acknowledgements

JL is funded by a Medical Research Council Scholarship (Award no. 1964851). SJT is funded by grants from the Bill & Melinda Gates Foundation (OPP1104516), the Biotechnology and Biological Sciences Research Council, the Department for International Development, The Economic and Social Science Research Council, The Natural Environment Research Council and the Defence, Science and Technology Laboratory, under the Zoonosis and Emerging and Livestock Systems (ZELS) programme (Grant no. BB/L019035/1). The Bill & Melinda Gates Foundation grant OPP1104516 also supports MCS. MCS acknowledges additional funding from the Medical Research Council (MR/M014975/1). The funders had no role in study design, data collection and analysis, decision to publish or preparation of the manuscript. The authors declare no conflicts of interest. The authors wish to thank Dr Simon Wagstaff and Mr Andrew Bennett for providing the computational resources to perform this analysis.

## Author contributions

SJT, MCS and JL conceived and planned the study. JL and AK were involved in field study design and data collection. JL wrote all computer code and designed and performed the analysis. JL wrote the first draft of the manuscript, and all authors contributed toward subsequent revisions. All authors gave final approval for publication.

## Data Availability

Due to file hosting limitations, both the 0.5m resolution and 3m resolution resistance surfaces are available upon request from the authors. Code used to generate and validate the resistance surfaces can be found as Supplementary File 2.

