## Supplementary File 1 for "Quantifying geographic accessibility to improve efficiency of entomological monitoring"

**This PDF file includes:**

Figs. S1 to S5

Table S1

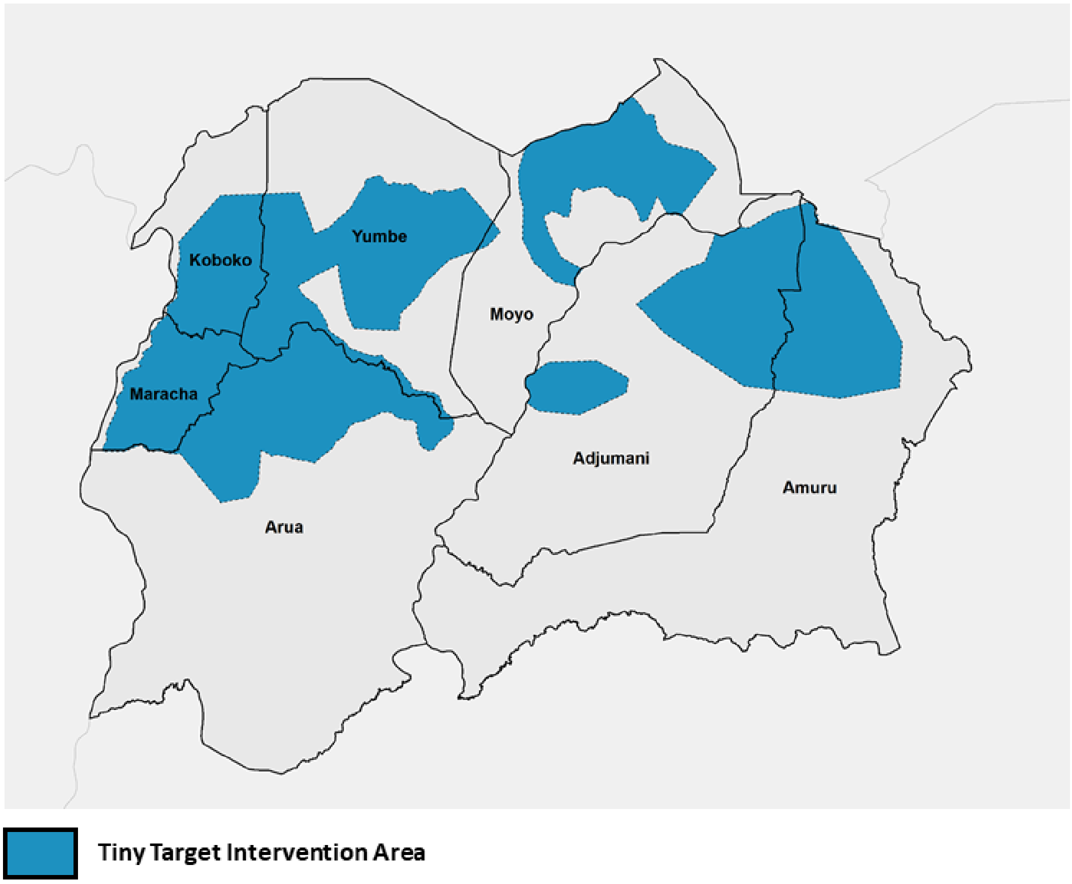


**Fig. S1. Existing and planned intervention areas.** Blue areas identify both current and planned Tiny Target intervention areas within the West Nile Region of Northern Uganda.

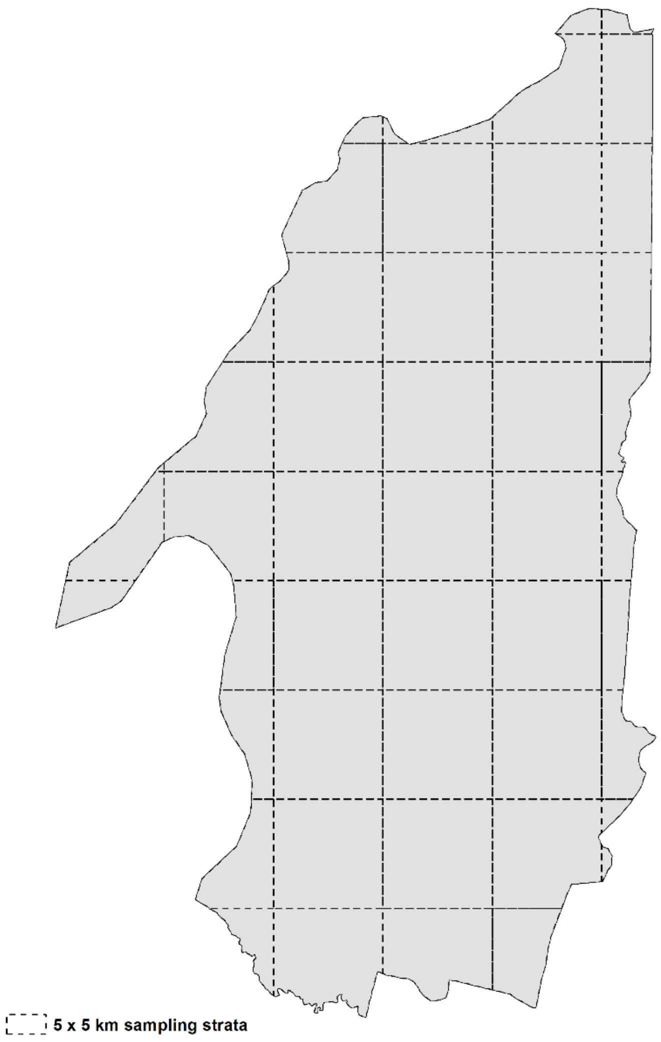


**Fig. S2. Distribution of 5 × 5 km sampling strata across Koboko district**.

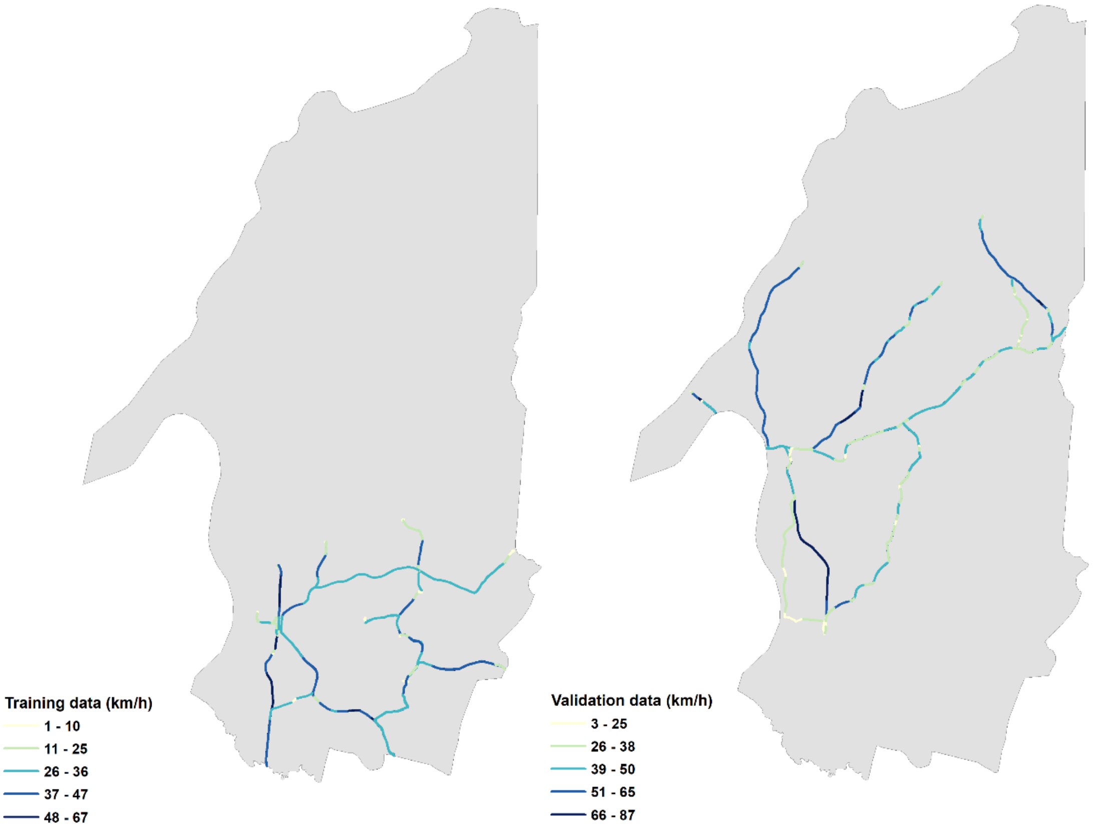


**Fig. S3. Spatial distribution of ground truth data.** Left: Tracks used to inform speeds along select roads (training data). Right: Tracks used to validate the generated cost-distance surface (validation data).

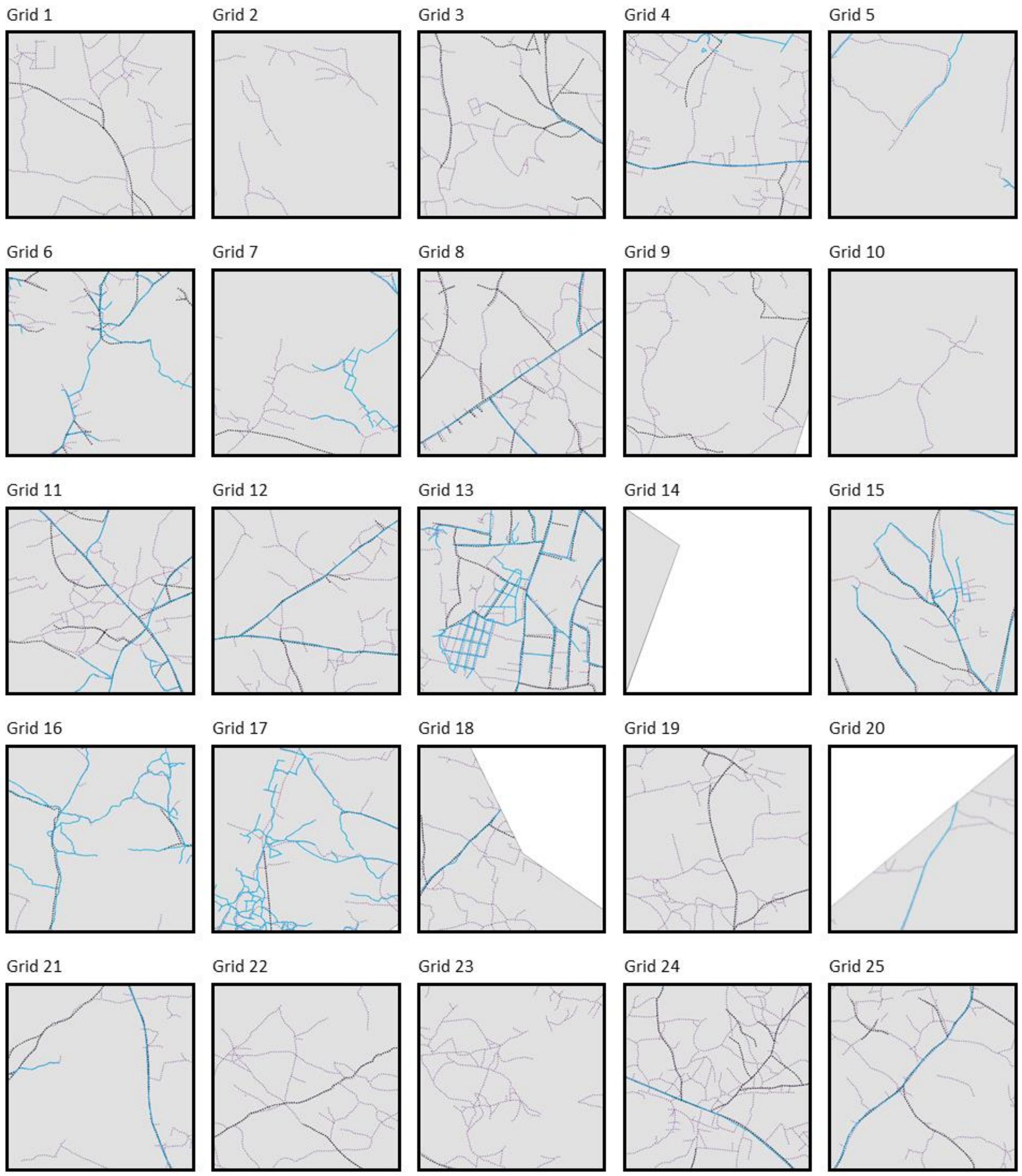


**Fig. S4. Composite images of digitised road networks within Koboko district.** Purple roads represent roads visible in 0.5m imagery; black roads represent roads visible in 3m imagery, and light blue roads represent roads available within the OSM dataset. The overlap of all three colours indicate areas of consistency across sources

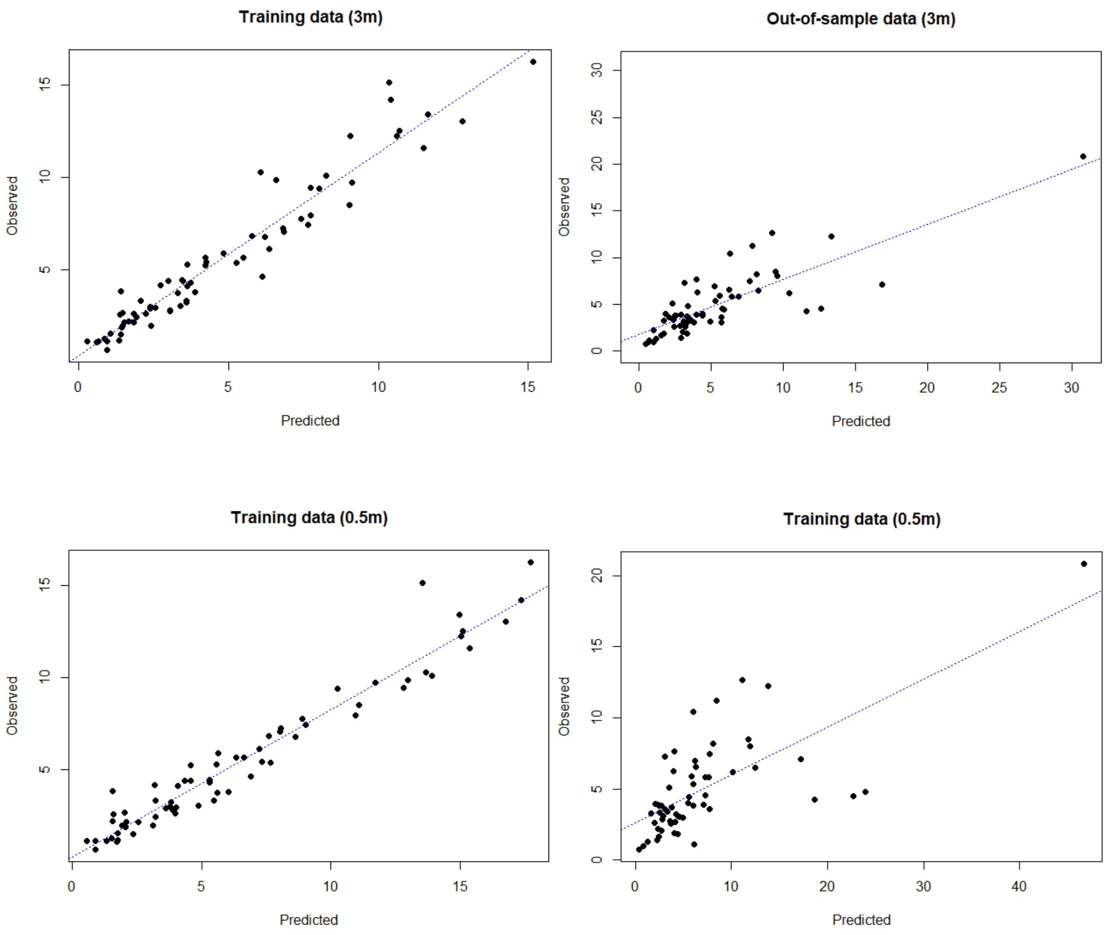


**Fig. S5. Regression plots.** Plots from a linear regression using observed travel time data with predicted travel time as the only covariate. Top Left: Regression using 3m within-sample (training) data. Top Right: Regression using 3m out-of-sample (validation) data. Bottom Left: Regression using 0.5m within-sample (training) data. Bottom Right: Regression using 0.5m out-of-sample (validation) data.

**Table S1. Results of OpenStreetMap data validation.**

| Grid | Length of road digitized (m) | | | Percentage similarity (%) | | |
|---|---|---|---|---|---|---|
| | 0.5m | 3m | OSM | OSM vs 0.5m | 3m vs 0.5m | OSM vs 3m |
| 1 | 7804.60 | 1459.33 | 0.00 | 0.00 | 18.70 | 0.00 |
| 2 | 3471.12 | 0.00 | 0.00 | 0.00 | 0.00 | 0.00 |
| 3 | 7759.08 | 3738.06 | 333.07 | 4.29 | 48.18 | 8.91 |
| 4 | 9146.50 | 1843.44 | 1817.94 | 19.88 | 20.15 | 98.62 |
| 5 | 2718.92 | 0.00 | 937.76 | 34.49 | 0.00 | 0.00 |
| 6 | 5515.98 | 2856.50 | 3483.76 | 63.16 | 51.79 | 121.96 |
| 7 | 3840.01 | 676.79 | 2254.52 | 58.71 | 17.62 | 333.12 |
| 8 | 10131.97 | 4607.29 | 1962.63 | 19.37 | 45.47 | 42.60 |
| 9 | 6053.58 | 1703.86 | 0.00 | 0.00 | 28.15 | 0.00 |
| 10 | 2034.04 | 0.00 | 0.00 | 0.00 | 0.00 | 0.00 |
| 11 | 12067.40 | 4540.82 | 3612.87 | 29.94 | 37.63 | 79.56 |
| 12 | 8940.11 | 2704.17 | 2082.35 | 23.29 | 30.25 | 77.01 |
| 13 | 14452.65 | 7053.10 | 9911.33 | 68.58 | 48.80 | 140.52 |
| 14 | 0.00 | 0.00 | 0.00 | 0.00 | 0.00 | 0.00 |
| 15 | 6650.08 | 3311.35 | 4432.14 | 66.65 | 49.79 | 133.85 |
| 16 | 3473.59 | 1370.35 | 5389.54 | 155.16 | 39.45 | 393.30 |
| 17 | 6779.36 | 772.51 | 7997.38 | 117.97 | 11.40 | 1035.25 |
| 18 | 5637.56 | 1214.75 | 649.64 | 11.52 | 21.55 | 53.48 |
| 19 | 9459.31 | 1912.29 | 0.00 | 0.00 | 20.22 | 0.00 |
| 20 | 2429.74 | 0.00 | 730.74 | 30.07 | 0.00 | 0.00 |
| 21 | 4846.82 | 2094.06 | 1343.11 | 27.71 | 43.20 | 64.14 |
| 22 | 8722.51 | 1564.75 | 0.00 | 0.00 | 17.94 | 0.00 |
| 23 | 8715.20 | 204.00 | 0.00 | 0.00 | 2.34 | 0.00 |
| 24 | 15891.30 | 4044.70 | 1064.19 | 6.70 | 25.45 | 26.31 |
| 25 | 8491.67 | 2918.87 | 1285.66 | 15.14 | 34.37 | 44.05 |
| **Total** | **175033.09** | **50590.99** | **49288.63** | **28.16** | **28.90** | **97.43** |
