## Supplementary File 2 for "Quantifying geographic accessibility to improve efficiency of entomological monitoring": 1.generate_classification_validation_points.R

```
# script to read in the maximum likelihood classified surface and generate random points to validate
# clear workspace
rm(list = ls())

#### load required packages ####
pacman::p_load(raster) # processing and manipulation of raster data

#### load in the classified raster, where this has already been processed (filtered, generalised etc.) ####
koboko <- raster("D:/Dropbox/PhD/1. Uganda/Accessibility/classification/0.5m classification/06FEB19/NIR_8BIT_50CM_SEG_MLE_FEBV2_025REJECT_filter_bc_rg_nibble.tif")

# values = 1 dirt road
# values = 2 tarmac road
# values = 3 dense vegetation
# values = 4 grassland
# values = 5 barren

## split to generate separate surfaces, one per class
## do this by retaining value and assigning all other cells the value of NA
dirt_roads <- koboko
m <- c(1.1, 9.1, NA)
road_rcm <- matrix(m, ncol = 3, byrow = TRUE)

beginCluster(8, type = "SOCK")
dirt_roads <- clusterR(dirt_roads, reclassify, args = list(rcl = road_rcm, right = FALSE))
endCluster()

tarmac_roads <- koboko
m <- c(0.9, 1.1, NA, 2.1, 9.1, NA)
road_rcm <- matrix(m, ncol = 3, byrow = TRUE)

beginCluster(8, type = "SOCK")
tarmac_roads <- clusterR(tarmac_roads, reclassify, args = list(rcl = road_rcm, right = FALSE))
endCluster()

dense_vegetation <- koboko
m <- c(0.9, 2.1, NA, 3.1, 9.1, NA)
road_rcm <- matrix(m, ncol = 3, byrow = TRUE)

beginCluster(8, type = "SOCK")
dense_vegetation <- clusterR(dense_vegetation, reclassify, args = list(rcl = road_rcm, right = FALSE))
endCluster()

grassland <- koboko
m <- c(0.9, 3.1, NA, 4.1, 9.1, NA)
road_rcm <- matrix(m, ncol = 3, byrow = TRUE)

beginCluster(8, type = "SOCK")
grassland <- clusterR(grassland, reclassify, args = list(rcl = road_rcm, right = FALSE))
endCluster()

barren <- koboko
m <- c(1.1, 4.1, NA, 5.1, 9.1, NA)
road_rcm <- matrix(m, ncol = 3, byrow = TRUE)

beginCluster(8, type = "SOCK")
barren <- clusterR(barren, reclassify, args = list(rcl = road_rcm, right = FALSE))
endCluster()

##### sample random points for validation ####
## using the binary surfaces, take a random sample of cells and retain these for using in the confusion matrix
dirt_points <- sampleRandom(dirt_roads, 10000, na.rm = TRUE, xy = TRUE)
dirt_points <- dirt_points[sample(100, replace = F),] 

tarmac_points <- sampleRandom(tarmac_roads, 100000, na.rm = TRUE, xy = TRUE)
tarmac_points <- tarmac_points[sample(100, replace = F),]

dense_vegetation_points <- sampleRandom(dense_vegetation, 10000, na.rm = TRUE, xy = TRUE)
dense_vegetation_points <- dense_vegetation_points[sample(100, replace = F),]

grassland_points <- sampleRandom(grassland, 10000, na.rm = TRUE, xy = TRUE)
grassland_points <- grassland_points[sample(100, replace = F),]

barren_points <- sampleRandom(barren, 10000, na.rm = TRUE, xy = TRUE)
barren_points <- barren_points[sample(100, replace = F),]

## merge together to create a dataframe with 500 points
aap <- rbind(dirt_points, tarmac_points, dense_vegetation_points, grassland_points, barren_points)
aap <- as.data.frame(aap)
names(aap) <- c("x", "y", "class")

## write these locations out as a csv, then use to perform a manual accuracy assessment and generate the confusion matrix
write.csv(aap,
          'D:/Dropbox/PhD/1. Uganda/Accessibility/classification/0.5m classification/Koboko_0.5m_SEG_5CLASS_MLE_FEB06_aap.csv',
          row.names = FALSE)
```
