## Supplementary File 2 for "Quantifying geographic accessibility to improve efficiency of entomological monitoring": 2.merge_sources_to_create_friction_surface.R

```
# clear workspace
rm(list = ls())

# set working directory
setwd("P:/accessibility")

# check if pacman is on the machine; if not, install then use to install/load other required packages
if (!require("pacman")) install.packages("pacman")

# load required packages
pacman::p_load(raster, rgdal, sp, maptools, rdrop2, rgeos, gdistance)

# load in dropbox token
# token <- readRDS("DropBox_token.rds")

# set temporary file directory
# check it's retained from last time; need to change path
write("TMPDIR = 'E:/temp'", file = file.path(Sys.getenv('R_USER'), '.Renviron'))
tempdir()

# create a raster which has the average of GPS polylines
# first load an extent raster
extent_raster <- raster("Koboko_Feb12.tif")

# load polylines shapefile
polylines <- shapefile('merged_segments_all.shp')

# convert polyline object to same projection as imagery
proj_ext <- proj4string(extent_raster)
polylines <- spTransform(polylines, proj_ext)

# rasterize and generate the mean
observed_speeds <- rasterize(polylines, extent_raster, "speed", fun = mean)

# write out averaged raster
writeRaster(observed_speeds,
            'output/observed_speeds',
            format = "GTiff",
            overwrite = TRUE)

# read in the raw OSM shapefile
osm_roads <- shapefile('OSM_Koboko.shp')

# project to same projection as the imagery
osm_roads <- spTransform(osm_roads, proj_ext)

# create speed column using average speeds from the GPS data
# osm_pst_roads <- osm_roads
# osm_pst_roads$speed <- 4.79
# osm_pst_roads$speed <- ifelse(osm_pst_roads$fclass == "primary", 100, osm_pst_roads$speed)
# osm_pst_roads$speed <- ifelse(osm_pst_roads$fclass == "secondary", 100, osm_pst_roads$speed)
# osm_pst_roads$speed <- ifelse(osm_pst_roads$fclass == "tertiary", 100, osm_pst_roads$speed)
# osm_pst_roads$speed <- ifelse(osm_pst_roads$fclass == "unclassified", 8.86, osm_pst_roads$speed)

osm_pst_roads <- osm_roads
osm_pst_roads$speed <- 4.79
osm_pst_roads$speed <- ifelse(osm_pst_roads$fclass == "primary", 56.98, osm_pst_roads$speed)
osm_pst_roads$speed <- ifelse(osm_pst_roads$fclass == "trunk", 56.98, osm_pst_roads$speed)
osm_pst_roads$speed <- ifelse(osm_pst_roads$fclass == "secondary", 55.06, osm_pst_roads$speed)
osm_pst_roads$speed <- ifelse(osm_pst_roads$fclass == "tertiary", 48.60, osm_pst_roads$speed)
osm_pst_roads$speed <- ifelse(osm_pst_roads$fclass == "unclassified", 37.70, osm_pst_roads$speed)
osm_pst_roads$speed <- ifelse(osm_pst_roads$fclass == "track", 8.86, osm_pst_roads$speed)
osm_pst_roads$speed <- ifelse(osm_pst_roads$fclass == "track_grade1", 8.86, osm_pst_roads$speed)
osm_pst_roads$speed <- ifelse(osm_pst_roads$fclass == "track_grade2", 8.86, osm_pst_roads$speed)

# rasterize
assummed_speeds <- rasterize(osm_pst_roads, extent_raster, "speed")

# write out averaged raster
writeRaster(assummed_speeds,
            'output/assummed_speeds_osm',
            format = "GTiff",
            overwrite = TRUE)

# read in the Ana's speeds assigned shapefile
osm_roads_ana <- shapefile('July_29_Slim_Trim_RoadsAndSpeeds.shp')

# project to same projection as the imagery
osm_roads_ana <- spTransform(osm_roads_ana, proj_ext)

# create speed column using averages of trackers
osm_roads_ana$average_speed <- ((osm_roads_ana$MARK_AVG)+(osm_roads_ana$MATT_AVG)+(osm_roads_ana$IG_AVG))/3

# rasterize and generate the mean
ana_speeds <- rasterize(osm_roads_ana, extent_raster, "average_speed")

# write out averaged raster
writeRaster(ana_speeds,
            'output/ana_speeds',
            format = "GTiff",
            overwrite = TRUE)

# now, convert from a speed to an associated travel time (cost)
# get speed in m/s, then account for distance
observed_speeds <- (4.24/((observed_speeds*1000)/3600))
# currently not enough disk space to process
assummed_speeds <- (4.24/((assummed_speeds*1000)/3600))
# currently not enough disk space to process
ana_speeds <- (4.24/((ana_speeds*1000)/3600))

# stack these, and keep in this priority order
# 1. observed, 2. ana, 3. assummed
merged <- merge(observed_speeds, ana_speeds)
merged <- merge(merged, assummed_speeds)

# read in the off-road surface
resistance <- raster("NDVI_Koboko.tif")

# define vector of resistance values for each NDVI value
m <- c(-1, 0.1, 200, 0.1, 0.2, 4.37, 0.2, 0.3, 6.14, 0.3, 1, 10.34)
rclmat <- matrix(m, ncol=3, byrow=TRUE)
resistance <- reclassify(resistance, rclmat)

# read in the classified imagery
classified <- raster('Koboko_8BIT_3M_5CLASS_JAN19_clip_bc_nibble.tif')

# read in the urban rural layer
ur_layer <- raster("koboko_urban_rural.tif")

# split by type
tarmac_roads <- classified
dirt_roads <- classified

ur_layer[ur_layer >0] <- 1

t_reclass_matrix <- matrix(c(-Inf, 1.1, 2.1, 1.1, 2.1, 10.1, NA, 1, NA), nrow = 3, ncol = 3)
r_reclass_matrix <- matrix(c(1.1, 10.1, NA), nrow = 1, ncol = 3)

message("Reclassifying tarmac values.")
beginCluster(8, type = "SOCK")
tarmac_roads <- clusterR(tarmac_roads, reclassify, args = list(rcl = t_reclass_matrix, right = FALSE))
endCluster()

message("Reclassifying dirt road values.")
beginCluster(8, type = "SOCK")
dirt_roads <- clusterR(dirt_roads, reclassify, args = list(rcl = r_reclass_matrix, right = FALSE))
endCluster()

ur_layer <- resample(ur_layer, dirt_roads)

dirt_urban <- mosaic(dirt_roads, ur_layer, fun = sum)

# create reclassification matrix
reclass_matrix <- matrix(c(-Inf, 1.8, 1.1, 2.1, NA, 0.30528), nrow = 2, ncol = 3)
# change cells less than 2 to NA and urban dirt cells to represent a speed of 50km/h
dirt_urban <- reclassify(dirt_urban, reclass_matrix, right = FALSE)

# rural dirt road cells can have the higher legal speed
dirt_roads <- reclassify(dirt_roads, cbind(0.9, 1.2, 0.19081), right = FALSE)

# mosaic the two speeds surfaces to have a separate time one
dirt_roads <- mosaic(dirt_roads, dirt_urban, fun = max)

# mosaic tarmac surface to highlight urban cells
tarmac_urban <- mosaic(tarmac_roads, ur_layer, fun = sum)

# create reclassification matrix
reclass_matrix <- matrix(c(-Inf, 1.8, 1.1, 2.1, NA, 0.30528), nrow = 2, ncol = 3)

# change cells less than 2 to NA and urban dirt cells to represent a speed of 50km/h
tarmac_urban <- reclassify(tarmac_urban, reclass_matrix, right = FALSE)

# rural dirt road cells can have the higher legal speed
tarmac_roads <- reclassify(tarmac_roads, cbind(0.9, 1.2, 0.15264), right = FALSE)

# mosaic the two speeds surfaces to have a separate time one
tarmac_roads <- mosaic(tarmac_roads, tarmac_urban, fun = max)

# merge classified tarmac/dirt
classified_speeds <- mosaic(dirt_roads, tarmac_roads, fun = min)

# write out classified road speeds
writeRaster(classified_speeds,
            'output/classified_speeds',
            format = "GTiff",
            overwrite = TRUE)

# now, merge road and off-road speeds
cost_surface <- merge(merged, classified_speeds)
cost_surface <- merge(cost_surface, resistance)

# change negative values to NA
cost_surface[cost_surface < 0] <- NA

# write out the cost raster
writeRaster(cost_surface,
            file = paste0('output/cost_surface_3m_', Sys.Date()),
            format = 'GTiff',
            overwrite = TRUE)

##### 0.5m work ######
rm(list = ls())
invisible(gc())

# read in the resistance layer
# read in the off-road surface
extent_raster <- raster("Koboko_50cm_imagery.tif")
gps_raster <- raster('output/observed_speeds.tif')

# convert gps_raster object to same projection as imagery
proj_ext <- proj4string(extent_raster)
proj_gps <- proj4string(gps_raster)

if(proj_ext != proj_gps){
  
  stop("Check the projections of the track data and the imagery.")
  
} else {
  
  # resample so that the gps raster is at a 0.5m resolution
  # first crop it to the extent of the area
  gps_raster_c <- crop(gps_raster, extent_raster)
  
  # inform progress
  message("Resampling the observed speeds.")
  
  # then resample so that it's at a 50cm resolution
  gps_raster_rs <- resample(gps_raster_c, extent_raster, method = "bilinear")
  
}

# create observed speeds raster
observed_speeds <- gps_raster_rs

# read in the raw OSM shapefile
osm_roads <- shapefile('OSM_Koboko.shp')

# project to same projection as the imagery
osm_roads <- spTransform(osm_roads, proj_ext)

# filter to only select primary, secondary, tertiary and unclassified
osm_pst_roads <- osm_roads

# create speed column using national speed limits
osm_pst_roads$speed <- 4.79
osm_pst_roads$speed <- ifelse(osm_pst_roads$fclass == "primary", 56.98, osm_pst_roads$speed)
osm_pst_roads$speed <- ifelse(osm_pst_roads$fclass == "trunk", 56.98, osm_pst_roads$speed)
osm_pst_roads$speed <- ifelse(osm_pst_roads$fclass == "secondary", 55.06, osm_pst_roads$speed)
osm_pst_roads$speed <- ifelse(osm_pst_roads$fclass == "tertiary", 48.60, osm_pst_roads$speed)
osm_pst_roads$speed <- ifelse(osm_pst_roads$fclass == "unclassified", 37.70, osm_pst_roads$speed)
osm_pst_roads$speed <- ifelse(osm_pst_roads$fclass == "track", 8.86, osm_pst_roads$speed)
osm_pst_roads$speed <- ifelse(osm_pst_roads$fclass == "track_grade1", 8.86, osm_pst_roads$speed)
osm_pst_roads$speed <- ifelse(osm_pst_roads$fclass == "track_grade2", 8.86, osm_pst_roads$speed)

# rasterize, first to 3m resolution, then resample to 0.5m (theory behind this is
# that I don't want all roads to only be one pixel wide, when they may be wider in reality)
# inform progress
message("Rasterizing OSM data: 0.5m.")
assummed_speeds <- rasterize(osm_pst_roads, gps_raster_c, "speed")

message("Resampling OSM raster: 50cm.")
assummed_speeds <- resample(assummed_speeds, extent_raster)

# write out averaged raster
message("Saving OSM raster: 50cm.")
writeRaster(assummed_speeds,
            file = 'output/koboko_assummed_speeds_osm_50cm',
            format = "GTiff",
            overwrite = TRUE)

# read in the Ana's speeds assigned shapefile
osm_roads_ana <- shapefile('July_29_Slim_Trim_RoadsAndSpeeds.shp')

# project to same projection as the imagery
osm_roads_ana <- spTransform(osm_roads_ana, proj_ext)

# create speed column using averages of trackers
osm_roads_ana$average_speed <- ((osm_roads_ana$MARK_AVG)+(osm_roads_ana$MATT_AVG)+(osm_roads_ana$IG_AVG))/3

# crop this to be the same extent as the district
osm_roads_ana <- crop(osm_roads_ana, extent_raster)

# rasterize at 3m, and generate the mean
message("Rasterizing Ana data: 0.5m.")
ana_speeds <- rasterize(osm_roads_ana, gps_raster_c, "average_speed")

message("Resampling Ana data: 50cm.")
ana_speeds <- resample(ana_speeds, extent_raster)

invisible(gc())

# write out averaged raster
writeRaster(ana_speeds,
            file = 'output/koboko_ana_speeds_50cm',
            format = "GTiff",
            overwrite = TRUE)

# now, convert from a speed to an associated travel time (cost)
# get speed in m/s, then account for distance
message("Converting observed speeds to times.")
observed_speeds <- (0.71/((observed_speeds*1000)/3600))

message("Converting assummed speeds to times.")
assummed_speeds <- (0.71/((assummed_speeds*1000)/3600))

message("Converting Ana's speeds to times.")
ana_speeds <- (0.71/((ana_speeds*1000)/3600))

# now, stack these, and keep in this priority
# 1. observed, 2. ana, 3. assummed
message("Merging obverved and assummed data.")
merged <- merge(observed_speeds, ana_speeds, overlap = TRUE)
merged <- merge(merged, assummed_speeds, overlap = TRUE)

# write out averaged raster
writeRaster(merged,
            file = 'output/merged_nc_50cm',
            format = "GTiff",
            overwrite = TRUE)

resistance <- raster("NDVI_50cm_Koboko.tif")

# define vector of resistance values for each NDVI value
m <- c(-1, 0.1, 200, 0.1, 0.2, 0.73, 0.2, 0.3, 1.03, 0.3, 1, 1.72)
rclmat <- matrix(m, ncol = 3, byrow = TRUE)

message("Reclassifying NDVI data.")
beginCluster(8, type = "SOCK")
resistance <- clusterR(resistance, reclassify, args = list(rcl = rclmat, right = FALSE))
endCluster()

# read in the classified 
classified <- raster("NIR_8BIT_50CM_SEG_MLE_FEBV2_025REJECT_filter_bc_rg_nibble.tif")

# read in the urban rural layer
ur_layer <- raster("koboko_urban_rural.tif")

# resample ur_layer to be 50cm
message("Resampling urban/rural layer.")
ur_layer <- resample(ur_layer, resistance)

message("Starting classification assignment.")
tarmac_roads <- classified
dirt_roads <- classified
t_reclass_matrix <- matrix(c(-Inf, 1.1, 2.1, 1.1, 2.1, 10.1, NA, 1, NA), nrow = 3, ncol = 3)
r_reclass_matrix <- matrix(c(1.1, 10.1, NA), nrow = 1, ncol = 3)

message("Reclassifying tarmac values.")
beginCluster(8, type = "SOCK")
tarmac_roads <- clusterR(tarmac_roads, reclassify, args = list(rcl = t_reclass_matrix, right = FALSE))
endCluster()

message("Reclassifying dirt road values.")
beginCluster(8, type = "SOCK")
dirt_roads <- clusterR(dirt_roads, reclassify, args = list(rcl = r_reclass_matrix, right = FALSE))
endCluster()

message("Reclassifying urban/rural values.")
beginCluster(8, type = "SOCK")
ur_layer <- clusterR(ur_layer, reclassify, args = list(rcl = cbind(0, 1000, 1), right = FALSE))
endCluster()

dirt_urban <- mosaic(dirt_roads, ur_layer, fun = sum)

# create reclassification matrix
reclass_matrix <- matrix(c(-Inf, 1.9, 1.9, 2.1, NA, 0.05112), nrow = 2, ncol = 3)

# change cells less than 2 to NA and urban dirt cells to represent a speed of 50km/h
message("Assigning urban dirt road travel times.")
beginCluster(8, type = "SOCK")
dirt_urban <- clusterR(dirt_urban, reclassify, args = list(rcl = reclass_matrix, right = FALSE))
endCluster()

# rural dirt road cells can have the higher legal speed
message("Assigning rural dirt road travel times.")
beginCluster(8, type = "SOCK")
dirt_roads <- clusterR(dirt_roads, reclassify, args = list(rcl = cbind(0.9, 1.2, 0.03651429), right = FALSE))
endCluster()

invisible(gc())

# mosaic the two speeds surfaces to have a separate time one
message("Combining dirt road travel times.")
dirt_roads <- mosaic(dirt_roads, dirt_urban, fun = max)

# mosaic tarmac surface to highlight urban cells
tarmac_urban <- mosaic(tarmac_roads, ur_layer, fun = sum)

# create reclassification matrix
reclass_matrix <- matrix(c(-Inf, 2.1, 2.1, 3.1, NA, 0.05112), nrow = 2, ncol = 3)

# change cells less than 2 to NA and urban tarmac cells to represent a speed of 50km/h
message("Reclassifying tarmac roads.")
beginCluster(8, type = "SOCK")
tarmac_urban <- clusterR(tarmac_urban, reclassify, args = list(rcl = reclass_matrix, right = FALSE))
endCluster()

# rural tarmac road cells can have the higher legal speed
beginCluster(8, type = "SOCK")
tarmac_roads <- clusterR(tarmac_roads, reclassify, args = list(rcl = cbind(1.9, 2.2, 0.02556), right = FALSE))
endCluster()

# mosaic the two speeds surfaces to have a separate time one
tarmac_roads <- mosaic(tarmac_roads, tarmac_urban, fun = max)

# merge classified tarmac/dirt
classified_speeds <- mosaic(dirt_roads, tarmac_roads, fun = min)

# now, merge road and off-road speeds
cost_surface <- merge(merged, classified_speeds)
cost_surface <- merge(cost_surface, resistance)

# write out the cost raster
writeRaster(cost_surface,
            file = paste0('output/cost_surface_50cm_', Sys.Date()),
            format = 'GTiff',
            overwrite = TRUE)
```
