## Supplementary File 2 for "Quantifying geographic accessibility to improve efficiency of entomological monitoring": 3.validation_linear_model.R

```
# clear workspace
rm(list = ls())

# read in the observed and predicted
data <- read.csv("D:/Dropbox/PhD/1. Uganda/Accessibility/resistance_validation/training_data_validation_2019-03-04_3m.csv",
                 stringsAsFactors = FALSE)

# generate a linear model between observed and pred
mod_3m <- lm(observed ~ predicted, data = data)

summary(mod_3m)
# Adjusted R-squared:  0.9292 

# residual sum of squares
RSS <- c(crossprod(mod_3m$residuals))

# mean squared error
MSE <- RSS/length(mod_3m$residuals)

# root MSE
RMSE <- RSS/mod_3m$df.residual

# read in the out-of-sample observed and predicted
data_oos <- read.csv("D:/Dropbox/PhD/1. Uganda/Accessibility/resistance_validation/outofsample_data_validation_2019-03-05_3m.csv",
                 stringsAsFactors = FALSE)

# generate a linear model between observed and pred
mod_oos_3m <- lm(observed ~ predicted, data = data_oos)

summary(mod_oos_3m)
# Adjusted R-squared:  0.6611 

# residual sum of squares
RSS_oos <- c(crossprod(mod_oos_3m$residuals))

# mean squared error
MSE_oos <- RSS_oos/length(mod_oos_3m$residuals)

# root MSE
RMSE_oos <- RSS_oos/mod_oos_3m$df.residual

# generate plot
plot(data$predicted, data$observed , pch = 16, col = "black",
     main = "Training data (3m)",
     xlab = "Predicted",
     ylab = "Observed")
abline(lm(observed ~ predicted, data = data), 
       col = "blue",
       lty = 3)

# generate plot
plot(data_oos$predicted, data_oos$observed , pch = 16, col = "black",
     main = "Out-of-sample data (3m)",
     xlab = "Predicted",
     ylab = "Observed",
     ylim = c(0, max(data_oos$predicted)),
     xlim = c(0, max(data_oos$predicted)))
abline(lm(observed ~ predicted, data = data_oos), 
       col = "blue",
       lty = 3)


# read in the observed and predicted for 0.5m (training)
data_50cm_training <- read.csv("D:/Dropbox/PhD/1. Uganda/Accessibility/resistance_validation/training_data_validation_2019-03-04_0.5m.csv",
                               stringsAsFactors = FALSE)

# generate a linear model between observed and pred
mod_50cm_training <- lm(observed ~ predicted, data = data_50cm_training)

summary(mod_50cm_training)
# Adjusted R-squared:  0.9462 

# residual sum of squares
RSS_50cm_training <- c(crossprod(mod_50cm_training$residuals))

# mean squared error
MSE_50cm_training <- RSS_50cm_training/length(mod_50cm_training$residuals)

# root MSE
RMSE_50cm_training <- RSS_50cm_training/mod_50cm_training$df.residual

# generate plot
plot(data_50cm_training$predicted, data_50cm_training$observed , pch = 16, col = "black",
     main = "Training data (0.5m)",
     xlab = "Predicted",
     ylab = "Observed")
abline(lm(observed ~ predicted, data = data_50cm_training), 
       col = "blue",
       lty = 3)

# read in the observed and predicted for 0.5m (OOS)
data_50cm_oos <- read.csv("D:/Dropbox/PhD/1. Uganda/Accessibility/resistance_validation/outofsample_data_validation_2019-03-05_0.5m.csv",
                               stringsAsFactors = FALSE)

# generate a linear model between observed and pred
mod_50cm_oos <- lm(observed ~ predicted, data = data_50cm_oos)

summary(mod_50cm_oos)
# Adjusted R-squared:  0.4883  

# residual sum of squares
RSS_50cm_oos <- c(crossprod(mod_50cm_oos$residuals))

# mean squared error
MSE_50cm_oos <- RSS_50cm_oos/length(mod_50cm_oos$residuals)

# root MSE
RMSE_50cm_oos <- RSS_50cm_oos/mod_50cm_oos$df.residual

# generate plot
plot(data_50cm_oos$predicted, data_50cm_oos$observed , pch = 16, col = "black",
     main = "Training data (0.5m)",
     xlab = "Predicted",
     ylab = "Observed")
abline(lm(observed ~ predicted, data = data_50cm_oos), 
       col = "blue",
       lty = 3)
```
