## Supplementary File 2 for "Quantifying geographic accessibility to improve efficiency of entomological monitoring": 4.run_cost_distance_algorithm.R

```
#### try and implement the cost-distance analysis ####
# clear the workspace and do a gc
rm(list = ls())
invisible(gc())

# set working directory
setwd("P:/accessibility")

# check if pacman is on the machine; if not, install then use to install/load other required packages
if (!require("pacman")) install.packages("pacman")

# load required packages
pacman::p_load(raster, rgdal, sp, maptools, rdrop2, rgeos, gdistance)

# set temporary file directory
# check it's retained from last time; need to change path
write("TMPDIR = 'E:/temp'", file = file.path(Sys.getenv('R_USER'), '.Renviron'))
tempdir()

#### start with the 3m ####
cost_3m <- raster('output/cost_surface_3m_2019-11-07.tif')

# load source location
source_point <- shapefile('Koboko_origin.shp')

# transform to UTM36N
proj_ext <- "+proj=utm +zone=36 +datum=WGS84 +units=m +no_defs +ellps=WGS84 +towgs84=0,0,0"
source_point <- spTransform(source_point, proj_ext)

# convert cost into a transition matrix
cost_transition <- transition(cost_3m, function(x) 1/mean(x), directions = 4)

# generate the cost surface
cost_distance_3m <- gdistance::accCost(cost_transition, source_point)

# convert to minutes
cost_distance_3m_minutes <- round(cost_distance_3m/60, 2)

# write out the difference surface
writeRaster(cost_distance_3m_minutes,
            "output/cost_distance_3m_minutes",
            format = "GTiff",
            overwrite = TRUE)

#### now run the 0.5m cd ####
# clear the workspace and do a gc
rm(list = ls())
invisible(gc())

cost_50cm <- raster('output/cost_surface_50cm_2019-11-07.tif') 

# load source location
source_point <- shapefile('Koboko_origin.shp')

# transform to UTM36N
proj_ext <- "+proj=utm +zone=36 +datum=WGS84 +units=m +no_defs +ellps=WGS84 +towgs84=0,0,0"
source_point <- spTransform(source_point, proj_ext)

# convert cost into a transition matrix
cost_transition <- transition(cost_50cm, function(x) 1/mean(x), directions = 4)

# generate the cost surface
cost_distance_50cm <- gdistance::accCost(cost_transition, source_point)

# convert to minutes
cost_distance_50cm_minutes <- round(cost_distance_50cm/60, 2)

# write out the difference surface
writeRaster(cost_distance_50cm_minute,
            "output/cost_distance_50cm_minutes",
            format = "GTiff",
            overwrite = TRUE)
```
