## Supplementary File 2 for "Quantifying geographic accessibility to improve efficiency of entomological monitoring": 5.identify_sentinel_sites.R

```
# script to identify optimal sentinel site placement
# clear workspace
rm(list = ls())

# set working directory
setwd('D:/Dropbox/PhD/1. Uganda/Accessibility/')

# load required packages
pacman::p_load(raster, gdistance)

# load in the cost surface (minutes)
cost_surface <- raster("cost_output/Koboko_3m_cost_distance_15022019_minutes.tif")

# load in the tsetse abundance surface 
tsetse_abundance <- raster("tsetse abundance/Pred_counts_josh_clipped_koboko.tif")

# read in 50m cell raster
grid_50m <- raster("tsetse abundance/50m_grid_sample.tif")

# read in the strata raster
stratum <- raster('tsetse abundance/5km_raster.tif')

# crop the abundance surface, and resample to same grid size as cost surface
tsetse_crop <- crop(tsetse_abundance, cost_surface)
tsetse_resampled <- resample(tsetse_crop, cost_surface, method = "bilinear")

# mask surface by the cost surface to remove cells outside of Koboko
tsetse_masked <- mask(tsetse_resampled, cost_surface)

# make a selection of cells which have the top 1% of abundance values
tsetse_df <- as.data.frame(tsetse_masked, xy = TRUE)
tsetse_df <- tsetse_df[!is.na(tsetse_df$Pred_counts_josh_clipped_koboko), ]
names(tsetse_df)[3] <- "abundance"

# get an index of which stratum cells are in
full_locs <- tsetse_df[c('x', 'y')]
full_locs <- SpatialPoints(full_locs, proj4string = CRS("+proj=utm +zone=36 +datum=WGS84 +units=m +no_defs +ellps=WGS84 +towgs84=0,0,0"))
strata_values <- extract(stratum, full_locs)
tsetse_df$strata <- strata_values

tsetse_df <- tsetse_df[complete.cases(tsetse_df), ]

loop_list <- unique(tsetse_df$strata)
loop_list <- loop_list[!is.na(loop_list)]

# loop through each strata
for(i in 1:length(loop_list)){
  
  j <- loop_list[i]
  
  # subset to just get records for that particular strata
  strata_i <- tsetse_df[tsetse_df$strata == j, ]
  
  tsetse_ordered <- strata_i[order(strata_i$abundance, decreasing = TRUE), ]

  # use these points to extract info on travel time from origin
  # transform the points to UTM 36N
  # first make it a spatialpoints object
  positions <- tsetse_ordered[c('x', 'y')]
  positions <- SpatialPoints(positions, proj4string = CRS("+proj=utm +zone=36 +datum=WGS84 +units=m +no_defs +ellps=WGS84 +towgs84=0,0,0"))
  
  # plot the coordinates on top of the surface, and extract cell values
  cost_values <- extract(cost_surface, positions)
  
  # merge this back on to the dataframe
  tsetse_ordered$cost <- cost_values
  
  # only select one point per 50m grid cell area
  # grab an index for a cell value based on lat long
  positions <- tsetse_ordered[c('x', 'y')]
  coordinates(positions) <- c("x", "y")
  proj4string(positions) <- CRS("+proj=utm +zone=36 +datum=WGS84 +units=m +no_defs +ellps=WGS84 +towgs84=0,0,0")
  positions <- spTransform(positions, CRS("+proj=longlat +datum=WGS84 +no_defs +ellps=WGS84 +towgs84=0,0,0"))
  
  # get duplicates (pixel number)
  tsetse_ordered$cell <- cellFromXY(grid_50m, positions)

  # remove duplicates
  tsetse_ordered <- tsetse_ordered[order(tsetse_ordered$abundance, tsetse_ordered$cost, decreasing = c(TRUE, FALSE)), ]
  deduplicated_points <- tsetse_ordered[!duplicated(tsetse_ordered$cell), ]
  
  # create separate ranking columns 
  deduplicated_points$ab_rank <- rank(-deduplicated_points$abundance)
  deduplicated_points$cost_rank <- rank(deduplicated_points$cost)
  deduplicated_points$combined_rank <- deduplicated_points$ab_rank + deduplicated_points$cost_rank
  
  # keep the X lowest ranked locations
  ten_locs <- deduplicated_points[order(deduplicated_points$combined_rank, decreasing = FALSE), ]
  
  n_per_strata <- 2
  
  if(nrow(ten_locs <= n_per_strata)){
  
    ten_locs <- ten_locs[1:n_per_strata, ]
  
  }
  
  # combine the estimates across all strata
  if(i == 1){
    
    combined <- ten_locs
    
  } else {
    
    combined <- rbind(combined,
                      ten_locs)
    
  }

}

combined <- combined[!is.na(combined$x),]

# write out the data frame with the 10 points per grid
write.csv(combined,
          'D:/Dropbox/PhD/1. Uganda/Accessibility/tsetse abundance/stratified_lowcost_high_abundance_locations_march_expanded.csv',
          row.names = FALSE)

min(combined$abundance)
max(combined$abundance)
mean(combined$abundance)

min(combined$cost)
max(combined$cost)
mean(combined$cost)

plot(cost_surface)
points(combined$x, combined$y, col = "blue", pch = 16)
```
