## Supplementary File 2 for "Quantifying geographic accessibility to improve efficiency of entomological monitoring": 6.run_travelling_salesperson_problem.R

```
# script to solve the travelling salesperson problem, using a transition matric and a set of points derived from 
# the previous analysis

# clear workspace
rm(list = ls())

# load required packages
pacman::p_load(gdistance, TSP, raster)

# load in the friction surface
# friction_surface <- raster("D:/Dropbox/PhD/1. Uganda/Accessibility/cost_output/cost_surface_3m_FEB19.tif")

# load in the locations to used in the TSP
locations <- read.csv("D:/Dropbox/PhD/1. Uganda/Accessibility/tsetse abundance/stratified_lowcost_high_abundance_locations_march_expanded.csv")

# check that points are in the same coordinate system
coordinates(locations) <- ~x+y

proj_ext <- "+proj=utm +zone=36 +datum=WGS84 +units=m +no_defs +ellps=WGS84 +towgs84=0,0,0"
proj4string(locations) <- proj_ext

# # convert the friction surface into a transition matrix
# friction_transition <- transition(friction_surface, transitionFunction = mean, directions = 4)
# 
# # save the transition matrix
# saveRDS(friction_transition, file = "D:/Dropbox/PhD/1. Uganda/Accessibility/cost_output/friction_transition_matrix_3m.rds")
friction_transition <- readRDS("D:/Dropbox/PhD/1. Uganda/Accessibility/cost_output/friction_transition_matrix_3m.rds")

# # add a gc to create some memory
# gc()

# create a distance matrix for the points
distance_matrix <- costDistance(friction_transition, locations, locations)
saveRDS(distance_matrix, file = "D:/Dropbox/PhD/1. Uganda/Accessibility/tsetse abundance/distance_matrix_3m_select_points.rds")
write.csv(distance_matrix,
          "D:/Dropbox/PhD/1. Uganda/Accessibility/tsetse abundance/distance_matrix_3m_select_points.csv",
          row.names = FALSE)

# implement the TSP
tsp <- TSP(distance_matrix)

route <- solve_TSP(tsp)

# create a polylines object which is the route
path_locations <- match(route, row.names(locations))
route_coordinates <- NULL
route_coordinates <- locations@coords[path_locations,]
names(route_coordinates) <- c("x", "y")#
route_coordinates <- as.data.frame(route_coordinates)
route_coordinates$id <- row.names(route_coordinates)

library(sp)
library(maptools)

points_to_line <- function(data, long, lat, id_field = NULL, sort_field = NULL) {
  
  # Convert to SpatialPointsDataFrame
  coordinates(data) <- c(long, lat)
  
  # If there is a sort field...
  if (!is.null(sort_field)) {
    if (!is.null(id_field)) {
      data <- data[order(data[[id_field]], data[[sort_field]]), ]
    } else {
      data <- data[order(data[[sort_field]]), ]
    }
  }
  
  # If there is only one path...
  if (is.null(id_field)) {
    
    lines <- SpatialLines(list(Lines(list(Line(data)), "id")))
    
    return(lines)
    
    # Now, if we have multiple lines...
  } else if (!is.null(id_field)) {  
    
    # Split into a list by ID field
    paths <- sp::split(data, data[[id_field]])
    
    sp_lines <- SpatialLines(list(Lines(list(Line(paths[[1]])), "line1")))
    
    # I like for loops, what can I say...
    for (p in 2:length(paths)) {
      id <- paste0("line", as.character(p))
      l <- SpatialLines(list(Lines(list(Line(paths[[p]])), id)))
      sp_lines <- spRbind(sp_lines, l)
    }
    
    return(sp_lines)
  }
}

# create a spatial lines object of the optimal route
route_lines <- points_to_line(route_coordinates, "x", "y")

# write out the optimal route
shapefile(route_lines,
          "D:/Dropbox/PhD/1. Uganda/Accessibility/tsetse abundance/travelling_salesman_selected_points_dec19_10.shp")
```
