## Supplementary File 2 for "Quantifying geographic accessibility to improve efficiency of entomological monitoring": README.txt

Order in which to process scripts, including a brief description of requirements:

1.generate_classification_validation_points
This script uses a classified raster, which is the output of the maximum likelihood classification algorithm, to generate a random sample of 500 points to use
for classification validation. The script creates separate surfaces, one per class, where the surface is binary. For this piece, we used five different classes
within the classification algorithm; therefore we generate five different surfaces and generate a random sample of 100 locations per surface. These points are
exported and are then used within a manual validation process, where the algorithm determined class is compared to the 'true' class. These data are then used to
create the confusion matrix.

Cells within the input raster must have the following values, representative of the following classes (as assigned through image classification in ArcGIS or QGIS, 
depending on software):
dirt road: "1"
tarmac road: "2"
dense vegetation: "3"
grassland: "4"
barren: "5"

2.merge_sources_to_create_friction_surface
This script uses a raster of classified roads (derived from image classification on remotely sensed imagery), a polygon/polyline dataset from OpenStreetMap, 
a NDVI surface derived from remotely sensed imagery, and ground truth GPS tracks to create a combined resistance surface.

3.validation_linear_model
Using the friction (resistance) surface, GPS tracks, and the 'least-cost-path' algorithm in QGIS, generate estimates of travel time for paired locations. Compile a csv
file where there's the observed (time taken from the GPS tracks) and the predicted (results from the least-cost-path). This script then runs a linear model to investigate
the relationship between observed and predicted travel times. This is used to evaluate the surface.

4.run_cost_distance_algorithm
This script used a shapefile/coordinates of the origin in Koboko, and the friction surface generated in a prior script. The accumulated (cumulative) cost of travel to 
ever cell within the surface is generated from the origin location. The output returned is the travel time in seconds; a section of the script converts this to minutes
to aid interpretability.

5.identify_sentinel_sites
Using the cost-distance output, a surface of tsetse abundance, sampling strata and a 50m x 50m cell raster, this script performs a cumulative ranking to identify the 
most efficient locations in which to place monitoring sites. 

6.run_travelling_salesperson_problem
Using the location of the sentinel sites, and the friction surface, this script implements Concorde's algorithm to solve the travelling salesperson problem and identify
the optimum order in which to visit the selected montioring sites.
